# Proteomic and phospho-proteomic longitudinal signatures of human skeletal muscle in lung cancer cachexia

**DOI:** 10.1101/2025.11.26.690647

**Authors:** Jonas Sørensen, Christian T. Voldstedlund, Edmund Battey, Zakarias Ogueboule, Andrea Irazoki, Johanne Modvig, Cathrine Knøs, Anna Hammershøi, Christian S. Carl, Steffen H. Raun, Oksana Dmytriyeva, Erik A. Richter, Seppo W. Langer, Lykke Sylow

## Abstract

Weight loss is a potentially deadly hallmark of many cancers, including lung cancer. In particular, the loss of skeletal muscle mass and function impairs survival and lowers quality of life. Despite being a major determinant of prognosis, the molecular drivers of muscle wasting remain ill-defined. Therefore, there is a critical need for human molecular data to support the development of effective therapies for this currently untreatable condition. Here, we utilize cutting-edge proteomics technology to longitudinally map the proteome and phosphoproteome of skeletal muscle from patients with newly diagnosed, advanced-stage non-small cell lung cancer during their treatment. Leveraging deep in vivo clinical phenotyping of activity, body composition, muscle quality, and nutritional risk, we identified 118/174 muscle proteins/phospho-sites associated with cachexia at diagnosis with indications of sexual dimorphism. Treatment altered 278 proteins and 1,155 phospho-sites, of which 137/91 proteins/phospho-sites were associated with muscle wasting. Our findings highlight disrupted calcium, anabolic, and stress signalling, alongside extracellular matrix and mitochondrial alterations, as key molecular features of cachexia in non-small cell lung cancer. These clinically anchored proteomic and phosphoproteomic signatures provide potential targets for future research.

## Main

Cancer accounts for nearly 10 million deaths annually worldwide, with lung cancer being the leading cause of cancer mortality^1^. Patients affected by lung cancer often experience an adverse loss of body weight known as cancer-associated cachexia (CAC) ^2–4^. Skeletal muscle wasting is particularly detrimental because it increases the risk of toxicity to anticancer treatment, decreases functional capacity, lowers quality of life, and reduces survival for the patients^5–7^. The lack of effective treatments for CAC places a significant burden on patients, families and society^8–10^. Thus, there is an urgent unmet need to identify treatment targets to stop muscle wasting in patients with CAC. Identifying such targets requires detailed molecular insights into human skeletal muscle protein regulation and posttranslational signaling at cancer diagnosis and during anticancer treatment^11^.

Mass spectrometry (MS)–based global and phospho-proteomics have emerged as powerful tools for unbiased, system-wide quantifications, including broad proteome alterations and focused analysis of posttranslational modification of specific proteins or pathways^12^. This approach has been leveraged in cancer research and diagnostics, supporting the utility of proteomics and phospho-proteomics in precision medicine and for drug target discovery^13,14^. Unravelling the global and phospho-proteomic changes in skeletal muscle from patients with cancer could provide important new insights into molecular changes and point to new therapeutic drug targets. Current molecular insights are derived mainly from mouse cachexia models^15–17^ but translation to human applications remains challenging. The reliance on non-translational pre-clinical models^18,19^ and limited availability of patient-derived molecular data^18,20^, significantly hampers therapeutic development and limits our understanding of the pathogenesis of cachexia.

Therefore, we undertook a comprehensive analysis of the muscle proteome and phospho-proteome in a representative cohort of patients with cancer to gain deeper insights into the pathogenesis of this complex condition. To this end, we performed global and phospho-proteomic analyses in muscles of patients with non-small cell lung cancer (NSCLC) at the time of diagnosis and after 12 weeks of first-line treatment to track molecular changes together with comprehensive clinical data, including body composition, clinical outcomes, functional measures, and patient-reported outcomes. We hypothesized that CAC present at NSCLC diagnosis and muscle wasting during first-line treatment are associated with distinct global and phospho-proteomic adaptations in skeletal muscle. By integrating these molecular changes with clinical data, we aimed to identify targetable signalling pathways underlying CAC and gain mechanistic insights into the pathophysiology of cachexia.

Integrating proteomic, clinical, and functional data, we identified 118 proteins and 174 phospho-sites linked to CAC at diagnosis. Treatment altered 278 proteins and 1,155 phospho-sites, of which 137/91 proteins/phospho-sites were associated with muscle wasting. These effects showed pronounced sexual dimorphism. Disrupted calcium signaling emerged as a central feature of CAC in non-small cell lung cancer and a potential therapeutic target.

## RESULTS

### Clinical characteristics of patients

To characterize the proteomic landscape of skeletal muscle in CAC and muscle wasting, paired biopsies of the vastus lateralis muscle were obtained from patients with advanced stage NSCLC; at diagnosis and after 12 weeks of first-line treatment (Fig. 1a). We included a total of 18 patients with advanced stage NSCLC. Mean age was 64 years, and there was an equal sex distribution. At the time of diagnosis, eight patients had CAC, four patients had pre-cachexia (preCAC), and six patients did not have cachexia (nonCAC). Fourteen patients received systemic intravenous treatment; six patients with immune check-point inhibitor, seven patients received platinum-based therapy in combination with immune check-point inhibitor, and one patient with vinorelbine. Four patients received oral tyrosine-kinase inhibitor as a tumor epidermal growth factor (EGFR) mutation was identified (Table 1). Eleven patients had Eastern Cooperative Oncology Group (ECOG) performance score 0, implying being fully able to carry out everyday activities. Overall, these characteristics were consistent with those reported in other NSCLC populations^21–23^. To better understand how body composition along with functional biochemical variables were associated with CAC at diagnosis we analyzed 18 clinical variables between patients with CAC and nonCAC including: sit to stand test, fitness level, hand grip strength, BMI, dual-energy X-ray absorptiometry (DXA)-derived adipose tissue mass (ATM), lean body mass (LBM), appendicular skeletal muscle index, computed tomography (CT)-derived skeletal muscle radio attenuation, skeletal muscle index, subcutaneous adipose tissue index, nutritional risk score, hemoglobin, percentage glycolated Hgb (HbA1c), albumin, c-reactive protein (CRP), and activity variables; time on feet, intensity count, step count (Fig. 1b). There was a 10% reduction in albumin in patients with CAC compared to nonCAC (Table S1). Of the atrogenes we observed a 57% increase in muscle MuRF1 but not Atrogin1 protein content in muscle from patients with CAC compared with nonCAC (Fig. 1c), corroborating preclinical observations on the activation of pro-atrophic molecular signatures in CAC^24^. In contrast, muscle fiber size distribution was similar between CAC and nonCAC groups (Fig. 1e+f).

**Figure 1.**
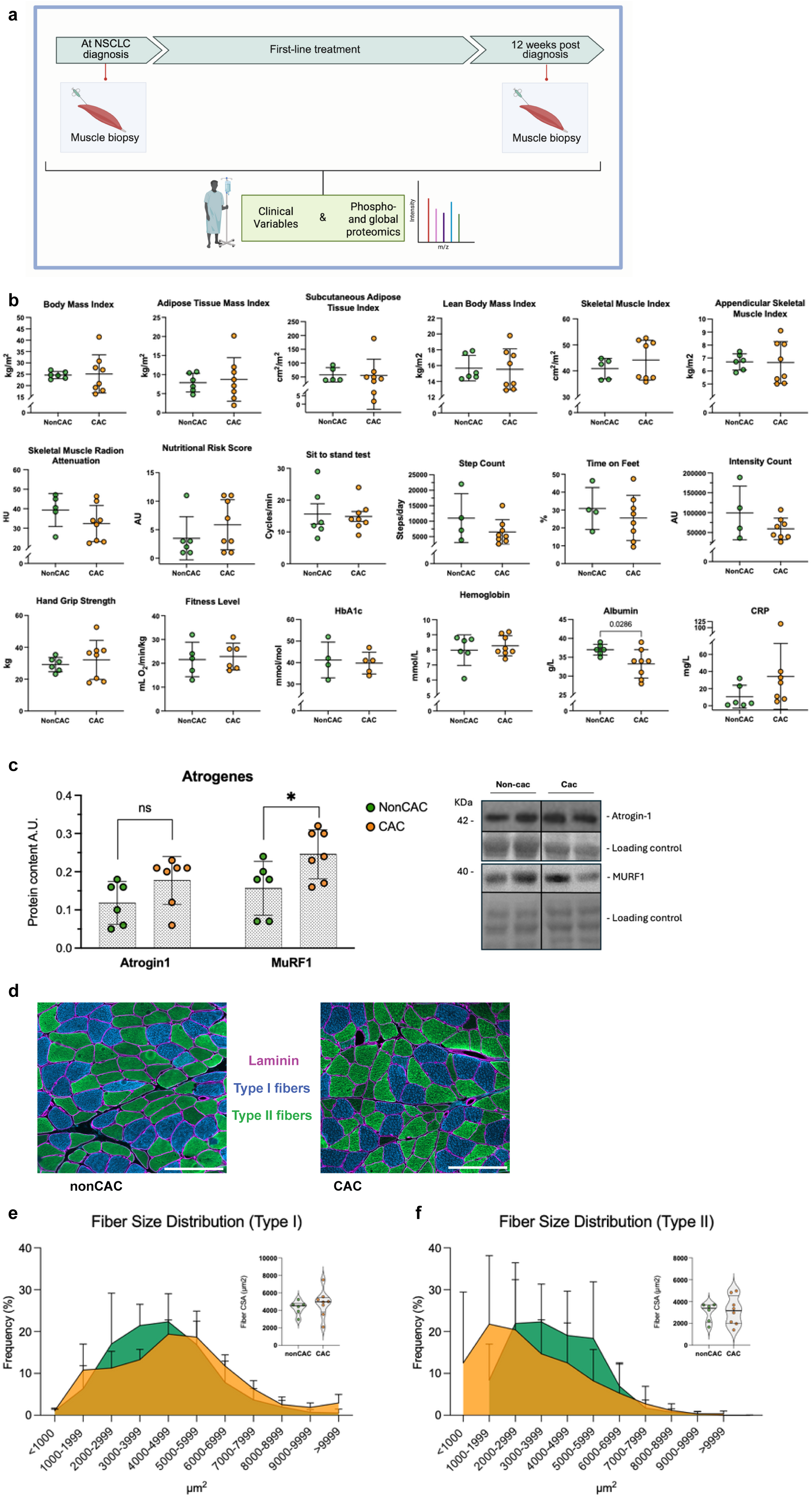
Assessment flow and baseline clinical and skeletal muscle phenotypes in patients with advanced stage NSCLC stratified by cachexia status **A:** Schematic representation of study workflow. Longitudinal data were obtained at diagnosis and week 12, including skeletal muscle biopsies and clinical assessment. Clinical variables along with phospho- and global proteomics were used for subsequent analyses. **B:** Comparison of clinical variables between nonCAC (green) and CAC (orange) patients. **C:** Protein expression levels of MuRF1 and Atrogin1 in skeletal muscle measured by western blotting at diagnosis. **D:** Representative images of cross-sectional area (CSA) of nonCAC patient (left) and CAC patient (right). Magenta: laminin, blue: type I fibers, green: type II fibers. Scale bars are 200 µm. **E:** Fiber size distribution and fiber cross-sectional area (CSA) of type I muscle fibers at diagnosis. **F:** Fiber size distribution and fiber cross-sectional area (CSA) of type II muscle fibers at diagnosis. Fiber size distribution data are presented as mean ± SE, while violin plot visualizes individual datapoints for fiber CSA. All other data are presented as mean ± SE with individual datapoints shown. Statistical comparisons were performed using Welch’s t-test. **Abbreviations:** MS, mass spectrometry; CRP, C-reactive protein; nonCAC, non-cachectic; CAC, cachectic; HgbA1c, glycated hemoglobin; AU, arbitrary units.

**Table 1.**
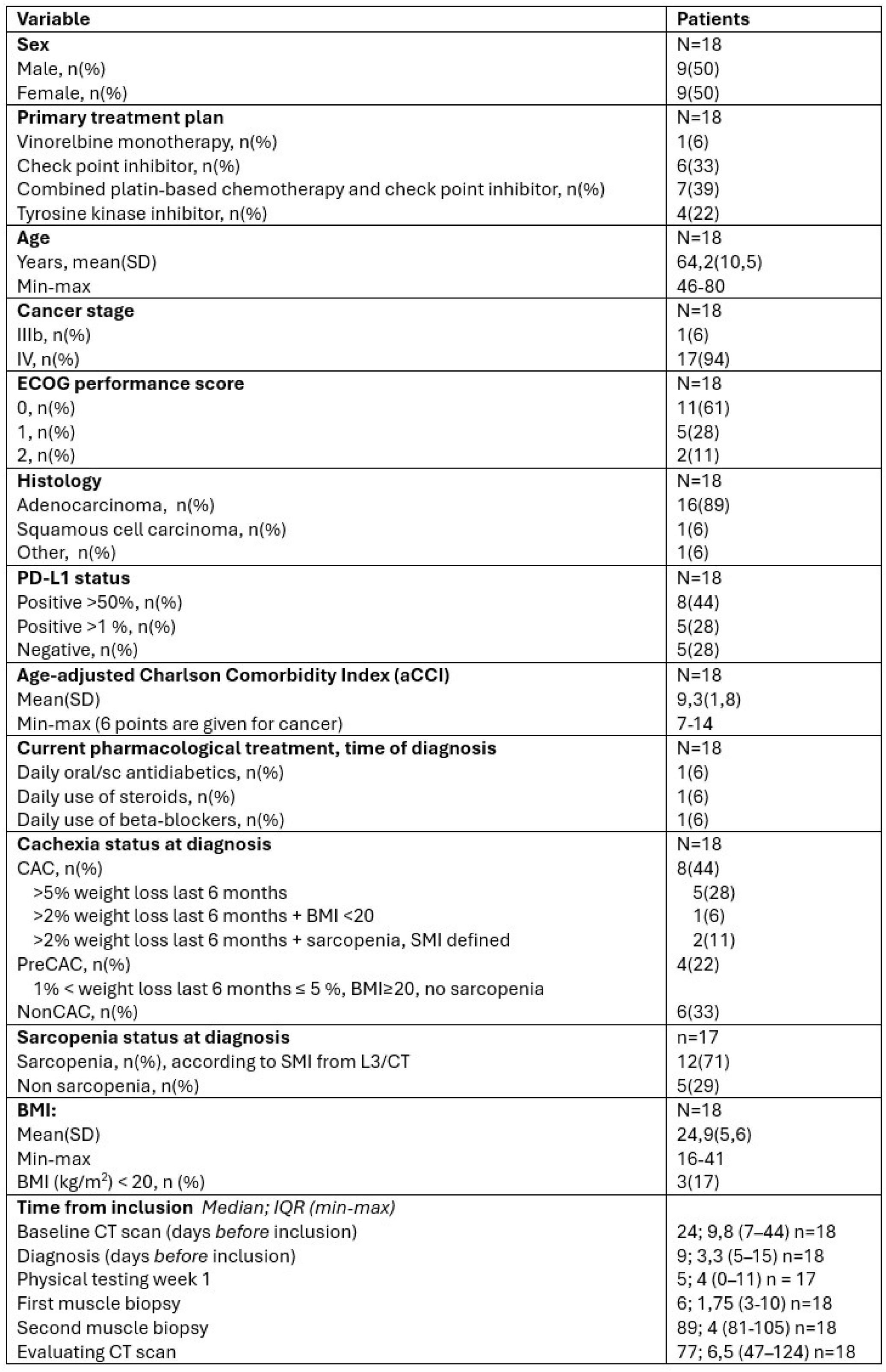
Patient characteristics at the time of diagnosis. Data are presented as *n* (%), mean (SD = standard deviation), unless otherwise specified. Percentages were calculated using the number of patients in each row as the denominator. **Abbreviations:** ECOG, Eastern Cooperative Oncology Group; PD-L1, programmed death-ligand 1; CAC, cachectic; PreCAC, pre-cachectic; NonCAC, non-cachectic; BMI, body-mass index; CT, computed tomography; SMI, skeletal muscle index; L3/CT, lumbar vertebral level 3 in computed tomography scan.

| **Variable** | **Patients** |
| --- | --- |
| **Sex** | N=18 |
| Male, n(%) | 9(50) |
| Female, n(%) | 9(50) |
| **Primary treatment plan** | N=18 |
| Vinorelbine monotherapy, n(%) | 1(6) |
| Check point inhibitor, n(%) | 6(33) |
| Combined platin-based chemotherapy and check point inhibitor, n(%) | 7(39) |
| Tyrosine kinase inhibitor, n(%) | 4(22) |
| **Age** | N=18 |
| Years, mean(SD) | 64,2(10,5) |
| Min-max | 46-80 |
| **Cancer stage** | N=18 |
| IIIb, n(%) | 1(6) |
| IV, n(%) | 17(94) |
| **ECOG performance score** | N=18 |
| 0, n(%) | 11(61) |
| 1, n(%) | 5(28) |
| 2, n(%) | 2(11) |
| **Histology** | N=18 |
| Adenocarcinoma, n(%) | 16(89) |
| Squamous cell carcinoma, n(%) | 1(6) |
| Other, n(%) | 1(6) |
| **PD-L1 status** | N=18 |
| Positive >50%, n(%) | 8(44) |
| Positive >1 %, n(%) | 5(28) |
| Negative, n(%) | 5(28) |
| **Age-adjusted Charlson Comorbidity Index (aCCI)** | N=18 |
| Mean(SD) | 9,3(1,8) |
| Min-max (6 points are given for cancer) | 7-14 |
| **Current pharmacological treatment, time of diagnosis** | N=18 |
| Daily oral/sc antidiabetics, n(%) | 1(6) |
| Daily use of steroids, n(%) | 1(6) |
| Daily use of beta-blockers, n(%) | 1(6) |
| **Cachexia status at diagnosis** | N=18 |
| CAC, n(%) | 8(44) |
| >5% weight loss last 6 months | 5(28) |
| >2% weight loss last 6 months + BMI <20 | 1(6) |
| >2% weight loss last 6 months + sarcopenia, SMI defined | 2(11) |
| PreCAC, n(%) | 4(22) |
| 1% < weight loss last 6 months ≤ 5 %, BMI ≥ 20, no sarcopenia |  |
| NonCAC, n(%) | 6(33) |
| **Sarcopenia status at diagnosis** | n=17 |
| Sarcopenia, n(%), according to SMI from L3/CT | 12(71) |
| Non sarcopenia, n(%) | 5(29) |
| **BMI:** | N=18 |
| Mean(SD) | 24,9(5,6) |
| Min-max | 16-41 |
| BMI (kg/m<sup>2</sup>) < 20, n (%) | 3(17) |
| **Time from inclusion** *Median; IQR (min-max)* |  |
| Baseline CT scan (days *before* inclusion) | 24; 9,8 (7–44) n=18 |
| Diagnosis (days *before* inclusion) | 9; 3,3 (5–15) n=18 |
| Physical testing week 1 | 5; 4 (0–11) n = 17 |
| First muscle biopsy | 6; 1,75 (3-10) n=18 |
| Second muscle biopsy | 89; 4 (81-105) n=18 |
| Evaluating CT scan | 77; 6,5 (47–124) n=18 |

### Global proteomics uncover CAC-associated distinctive pathophysiologies related to muscle calcium signaling in patients with NSCLC at diagnosis

Having established the CAC stratification, clinical variables, and atrophy markers; we next characterized the proteomic landscape of skeletal muscle in patients with CAC (4 females/4 males) and nonCAC (3 females/3 males) using MS–based global proteomics. To identify targets defining the CAC phenotype, the following analysis focused on the comparison between nonCAC and CAC, representing the most distinct phenotypic contrast (Fig. 2a). We quantified a total of 3511 proteins (Fig. 2b), with samples exhibiting a high degree of similarity across biological sex and cachectic status (nonCAC & CAC) (Fig. S1a-c). This was further evidenced by the observed coefficients of variance (Fig. S1b) of 2.3% and 2% in nonCAC and CAC, respectively. No obvious sample separation across principal component 1 or 2 was evident, indicating the effect of CAC on skeletal muscle presents in a targeted manner, involving a subset of protein complexes and structures as opposed to a global alteration in protein expression (Fig. S1a + S1c). These data revealed 73 upregulated and 45 downregulated muscle proteins in patients with CAC compared to nonCAC at diagnosis (Fig. 2b), further indicating that alterations in muscle protein of patients with CAC are highly selective.

**Figure 2.**
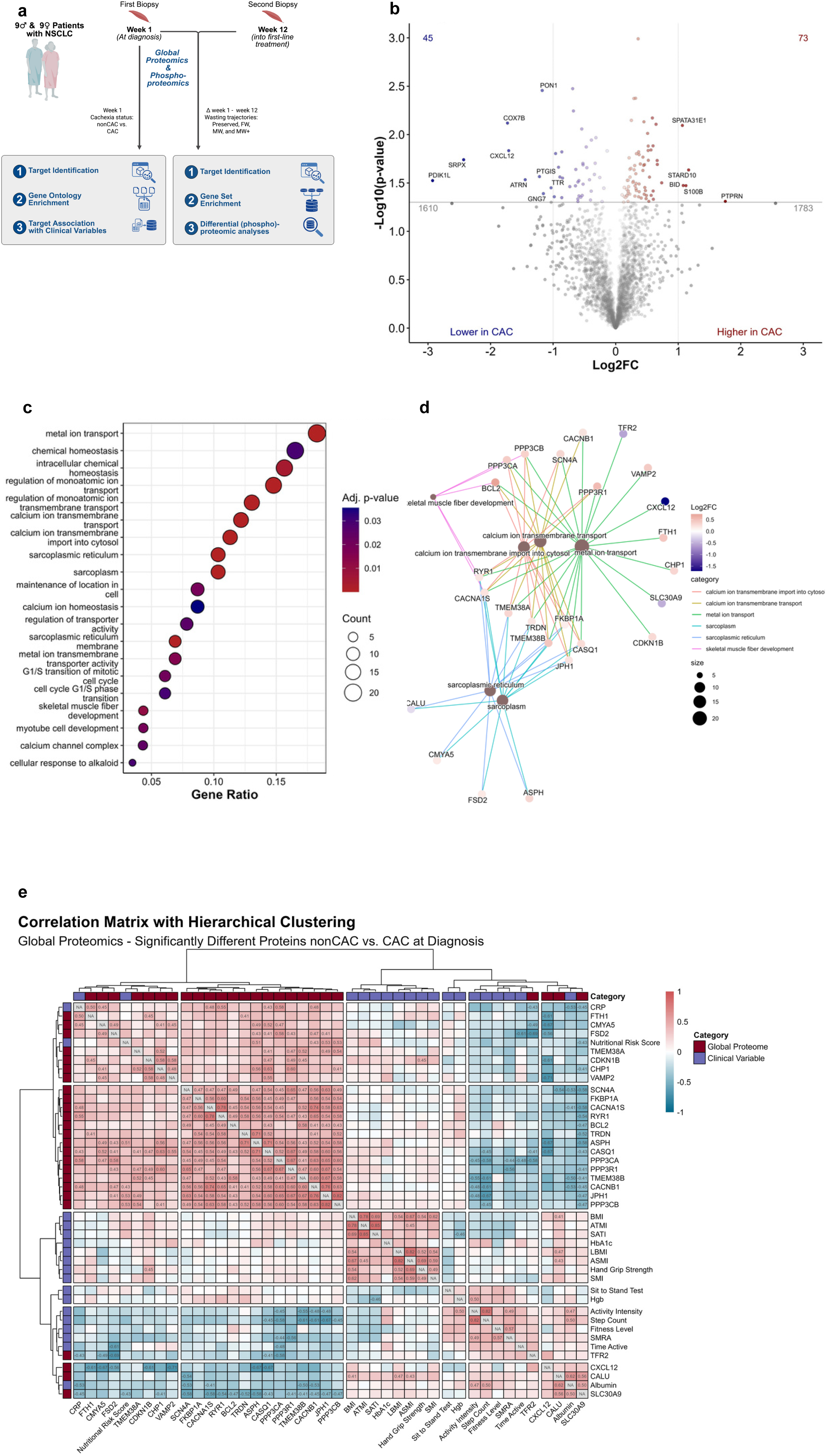
Global skeletal muscle proteomics in patients with NSCLC stratified by cachectic status identifies dysregulated calcium signaling **A:** Schematic representation of workflow and methods. 18 patients with non-small cell lung cancer (NSCLC) were included and 2 muscle biopsies were obtained at diagnosis (week 1) and 12 weeks later. Data derived from week one informed the stratification of cachectic (CAC) compared to non-cachectic (nonCAC) patients with subsequent analyses. Based on differences in clinical measures from week 1 to week 12, patients were stratified into wasting phenotypes; preserved, fat wasting (FW), muscle wasting (MW) and muscle wasting with additional weight loss (MW+). Analyses investigating the proteomic and phosphoproteomic changes from week 1 to week 12 were carried out. **B:** Volcano plot displaying the log2 fold change (Log2FC) and -log10 p-values of proteins quantified by global proteomics at diagnosis. Significantly regulated proteins with Log2FC > 1 or < −1 are highlighted. Statistical comparisons were performed using Welch’s t-test, and multiple-comparison adjustment was applied using permutation-based correction. **C:** Over-representation analysis of all significantly regulated proteins in CAC versus nonCAC at diagnosis. Gene Ontology (GO) categories included Cellular Component (CC), Biological Process (BP), and Molecular Function (MF). The top 20 significantly enriched GO terms are shown, ranked by gene ratio. P-values were corrected for multiple testing using the Benjamini–Hochberg false discovery rate (FDR) method **D:** Concept network showing proteins associated with the top five most significantly enriched GO terms, as well as the term *skeletal muscle fiber development*. Node size represents the number of proteins linked to each GO term and edge color representing term association. The corresponding Log2 fold change (Log2FC) for each protein in CAC versus nonCAC is indicated by color. **E:** Correlation matrix with hierarchical clustering illustrating associations between the 25 proteins identified in the concept network, clinical variables measured in the patients, and established markers of cancer cachexia identified in the same cohort in a separate study. Red indicates positive correlations, while blue indicates inverse correlations. Each box displays the Kendall correlation coefficient when the correlation was significant; otherwise, only the color indicates the direction of the association.

Next, we investigated whether proteins exhibiting different abundance between nonCAC and CAC patients were associated with cellular processes related to muscle function. Intriguingly, overrepresentation analysis (Fig. 2c) using Gene Ontology (GO) categories (cellular component, biological process, and molecular function) revealed enrichment of several processes related to ion transport, calcium homeostasis and skeletal muscle development, while the bulk regulation was localized to the sarcoplasmic reticulum. To further investigate the core regulatory components of cachexia, we extracted all proteins associated with the top 5 most significantly enriched GO terms as well as *“skeletal muscle fiber development”* (Fig. 2d), revealing a highly interconnected network of 25 proteins. Among these, 5 proteins (PPP3CA, PPP3CB, RYR1, BCL2 and CACNA1S) exhibited connectivity across both the most significantly regulated terms as well as *“skeletal muscle fiber development”*, potentially indicating a core regulatory role in calcium-dependent signaling in cancer cachexia.

### Key clinical variables and measures of body composition were correlated with calcium-regulated components of the global muscle proteome at diagnosis

To investigate the clinical relevance of the 25 proteins identified in the network analysis, we performed a correlation analysis including 18 clinical variables. Of the 25 CAC-related muscle proteins, 17 correlated significantly with key clinical variables (Fig. 2e) (CDKN1B, ASPH, CASQ1, PPP3CA, PPP3CB, PPP3R1, SCN4, TMEM38B, CACNB1, JPH1, CACNA1S, RYR1, FTH1, CMYA5, FSD2, CALU and SLC30A9).

Interestingly, the two calcineurin subunits, PPP3CA and PPP3CB, involved in calcium signaling, emerged as potential core signaling nodes, due to involvement in immune regulation and muscle development^25^. This was supported by significant positive correlations between PPP3CA and CRP (τ = 0.58) and a significant inverse correlation with step count (τ = –0.58), intensity count (τ = –0.45), and skeletal muscle radio attenuation (τ = –0.44) (Fig. 2e). In turn, PPP3CB was positively correlated with nutritional risk score (τ = 0.53) and inversely correlated with step count (τ = –0.45). Three additional calcium-related proteins correlated with CRP, namely CASQ1 (τ = 0.43), involved in calcium storage and release during contraction^26^, CMYA5 (τ = 0.45), critical for excitation-contraction coupling in both cardiac and skeletal muscles^27^, and CACNA1S (τ = 0.48), a calcium-channel subunit, involved in muscle contraction and hypermetabolism in the skeletal muscles when gene-mutated^28^. Lastly, CALU, a calcium-binding protein localized in the sarcoplasmic reticulum, and downregulated in CAC, showed positive correlation with appendicular skeletal muscle index (τ = 0.43), LBM (τ = 0.47), BMI (τ = 0.41), and albumin (τ = 0.62), while SLC30A9, a mitochondrial zinc transporter^29^ and downregulated in CAC, showed inverse correlation with nutritional risk score (τ = –0.43) and CRP (τ = –0.45) (Fig. 2e).

By correlating the muscle proteome with clinical variables at NSCLC diagnosis (Fig. 2e), we provide an first clinically anchored muscle biobank from a discovery cohort of NSCLC patients, which may serve as a foundation for future validation in larger cohorts. These data indicate selective alterations enriched in ion transport, calcium homeostasis, and sarcoplasmic reticulum biology, converging on a 25-protein network with calcineurin and calcium-channel components as central nodes. These protein-to-phenotype links highlight a coordinated altered calcium-signaling axis in CAC.

### Pathological post-translational modifications of phospho-proteins in skeletal muscle of NSCLC patients with CAC

After identifying global proteomic CAC-associated skeletal muscle alterations in patients with NSCLC at diagnosis, we determined post-translational alterations that could contribute to CAC at NSCLC diagnosis. As phosphorylation regulates protein activity^30^, we leveraged quantitative phospho-proteomics to explore signaling pathways and cellular processes involved in human CAC. We quantified 4,481 phosphorylated sites in muscle at NSCLC diagnosis, with 99 up- and 75 downregulated sites in CAC versus nonCAC patients (Fig. 3a). Compared to the proteome, the muscle phosphoproteome was overall less regulated in CAC, but exhibited greater inter-subject variability (Fig. S1b), suggesting a higher degree of heterogeneity at the phosphoproteome level in muscle of patients with NSCLC. Consistent with the global proteome (Fig. S1a,c), the phosphoproteome showed no clear separation across groups or sex (Fig. S2a,b), suggesting targeted CAC effects on skeletal muscle involving a subset of phosphoproteins.

**Figure 3.**
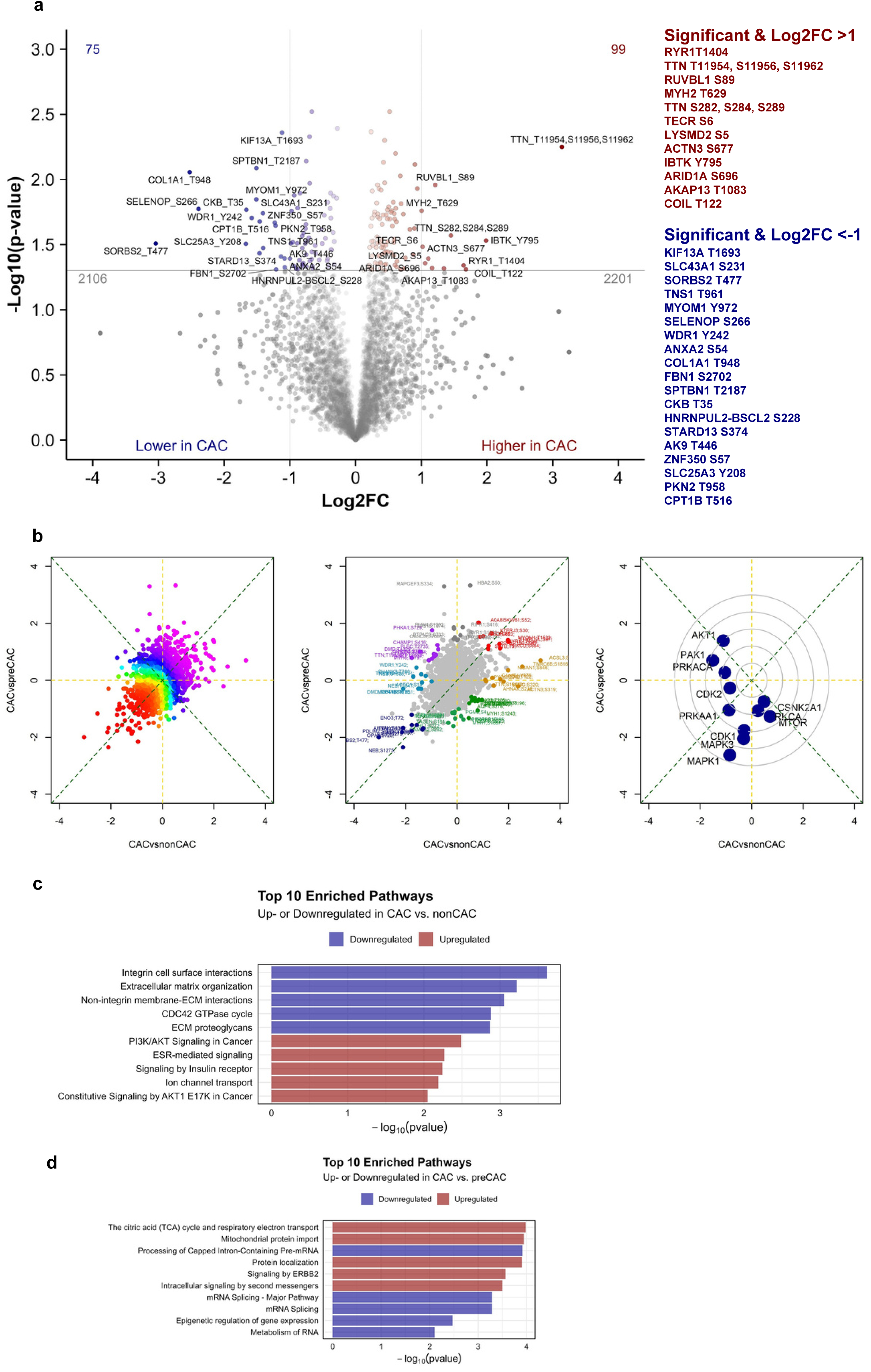
Phospho-proteomics reveals disrupted calcium, anabolic and stress signaling in the skeletal muscle of cachectic patients with NSCLC **A:** Volcano plot displaying the log2 fold change (Log2FC) and -log10 p-values of proteins quantified by phosphoproteomics at diagnosis. Significantly regulated proteins with Log2FC > 1 or < −1 are highlighted. Statistical comparisons were performed using Welch’s t-test, and multiple-comparison adjustment was applied using permutation-based correction. **B:** Kinase perturbation analysis. *Left:* Direction of regulation across the Log2 fold change (Log2FC) range for CAC versus nonCAC and CAC versus preCAC. *Middle:* Annotation of proteins across the Log2FC range for CAC versus nonCAC and CAC versus preCAC. *Right:* Kinase perturbation profiles inferred from the Log2FC distributions for CAC versus nonCAC and CAC versus preCAC. **C:** Gene-centric Reactome pathway enrichment analysis based on proteins quantified by phosphoproteomics at diagnosis in the comparison between CAC and nonCAC. The top five most significantly upregulated and top five most significantly downregulated pathways are shown. p-values were corrected for multiple testing using the Benjamini–Hochberg (BH) method. **D:** Gene-centric Reactome pathway enrichment analysis based on proteins quantified by phosphoproteomics at diagnosis in the comparison between CAC and preCAC. The top five most significantly upregulated and top five most significantly downregulated pathways are shown. p-values were corrected for multiple testing using the Benjamini–Hochberg (BH) method.

Among the 99 CAC upregulated phospho-sites, 12 sites were regulated with a Log2FC of >1.0 (Fig. 3a), while 19 of the 75 sites downregulated in CAC versus nonCAC patients, were regulated with a Log2FC of <-1.0. Overall, phosphorylation of sarcomeric (TTN, MYH2, ACTN3, RYR1), metabolic (CPT1B, CKB, TECR), and signaling/scaffold proteins (AKAP13, ARID1A, IBTK, COIL, RUVBL1) indicates coordinated regulation of muscle structure, energy metabolism, and calcium-dependent signaling pathways. Examining phosphosites significantly regulated in CAC versus non-CAC muscle that also showed significant differences in protein abundance, we identified 11 sites on 7 different proteins, of which 3 were composite (CKB T35; ABLIM2 S17; TECR S96; TECR S6; RYR1 T1404; CACNB1 S193; ASPH S97; ALPK3 S1390; JPH1 T461,S465; JPH1 T461,S465,S469 and JPH1 S171,S174).

Intriguingly, when subjecting these 11 sites to a correlation network analysis including the previously identified targets and variables (filtering for correlations with tau >0.6 or <-0.6, Fig. S2c), we observed several sites exhibiting strong linkage with the proteins associated with CAC in the current analysis. Although speculative, this may suggest that these sites are involved in mediating the molecular signaling related to CAC or represent compensatory modification to protein activity ameliorating the cellular perturbations induced by cancer.

Next, we performed a kinase-enrichment analysis to investigate whether any systematic patterns emerged, linking the phosphoproteome with the global proteome in relation to CAC. To delineate molecular features associated with the progression toward the CAC phenotype, the kinase perturbation analysis incorporated the preCAC group, enabling assessment of kinase activity changes across the phenotypic continuum from nonCAC to CAC. Intriguingly, we found 4 significantly perturbed kinases, namely; MAPK3, MAPK1, PRKAA1, and CDK1 (p < 0.05; Fig. 3b) (encoding Extracellular signal-Regulated kinase 1 and 2 (ERK1/2), 5’-AMP-activated protein kinase catalytic subunit alpha-1 (AMPKa1), and Cyclin-dependent Kinase 1 (CDK1), respectively). The perturbation of ERK1/2, AMPKa1, and CDK1 is suggestive of reduced kinase activity in CAC compared to non- and preCAC. In skeletal muscle, these four kinases are core integrators of energy and growth signaling^31–34^, and the collective downregulation indicates that CAC involves suppression of anabolic and metabolic control, consistent with the gradual CAC-related decline in skeletal muscle mass.

Subsequent gene-centric enrichment analysis revealed a prominent loss of ECM/adhesion and CDC42 signaling in CAC relative to nonCAC, alongside remodeling of ion channel and growth-factor pathways (Fig. 3c). Compared with preCAC, CAC showed enrichment within mitochondrial oxidative and import pathways but reduced RNA processing/splicing, consistent with stress adaptation under diminished anabolic capacity (Fig. 3d). Notably, insulin-receptor signaling was enriched in CAC versus nonCAC; however, in the context of reduced ECM/CDC42 signaling and suppressed ERK1/2 kinase signatures (Fig. 3b), this most likely represents a compensatory mechanism reflecting receptor-proximal phosphorylation rather than improved insulin action. Likewise, ERBB2 signaling was enriched in CAC relative to preCAC, indicating progressive growth-factor receptor activation, yet the concurrent reduction in RNA-processing capacity and anabolic kinase activity suggests a distal hampering in pathway functionality.

Together, these findings point to a decoupling of receptor signaling from mechano-transduction and energy-sensing circuits, compatible with impaired anabolic control and ongoing muscle loss in CAC and provide the first open-source muscle *CACProteome&Phosphoproteome* dataset clinically anchored to patient data.

### Longitudinal proteomic and phosphoproteomic alterations in skeletal muscle during first-line NSCLC treatment

Having identified the molecular signatures distinguishing patients with CAC and nonCAC at NSCLC diagnosis, we next investigated how skeletal muscle proteomes changed over 12 weeks of first-line NSCLC treatment for each individual patient. Leveraging our longitudinal design (Fig. 2a), which included paired muscle biopsies and clinical assessments from the same patients in week 1 and week 12, we were able to evaluate treatment-associated molecular and functional changes.

Muscle biopsied after 12 weeks of first-line NSCLC treatment exhibited multiple proteomic changes compared with muscle collected during the first week of diagnosis, with 152 proteins upregulated and 126 downregulated (Fig. 4a), and 647 upregulated and 508 downregulated phosphosites (Fig. 4b). The extent of these changes was further supported by principal component analysis, revealing distinct phosphoproteome clustering possibly reflecting treatment-dependent differences in phosphorylation patterns from time of diagnosis to week 12 (Fig. S3c,d). In contrast, the global proteome showed no clear separation (Fig. S3a,b), suggesting that post-translational phosphorylation is more dynamically regulated during NSCLC treatment than overall protein abundance. Further gene set enrichment analysis (GSEA) of the skeletal muscle proteome revealed marked activation of immune and inflammatory pathways after treatment (Fig. 4c), consistent with immune stimulation characteristic of immune checkpoint blockade received by 72% of patients. Complementing this, phosphoproteomic pathway enrichment revealed coordinated post-translational regulation of stress-responsive and transcriptional signaling networks, including FOXO-mediated cell death programs, EGFR and EPH-Ephrin signaling, unfolded protein response, and mitophagy (Fig. 4d). Given that most patients received immune checkpoint inhibitors, alone or in combination with platin-based chemotherapy, the integrated proteome–phosphoproteome response points to a treatment-driven immune–metabolic activation that has not been previously characterized.

**Figure 4.**
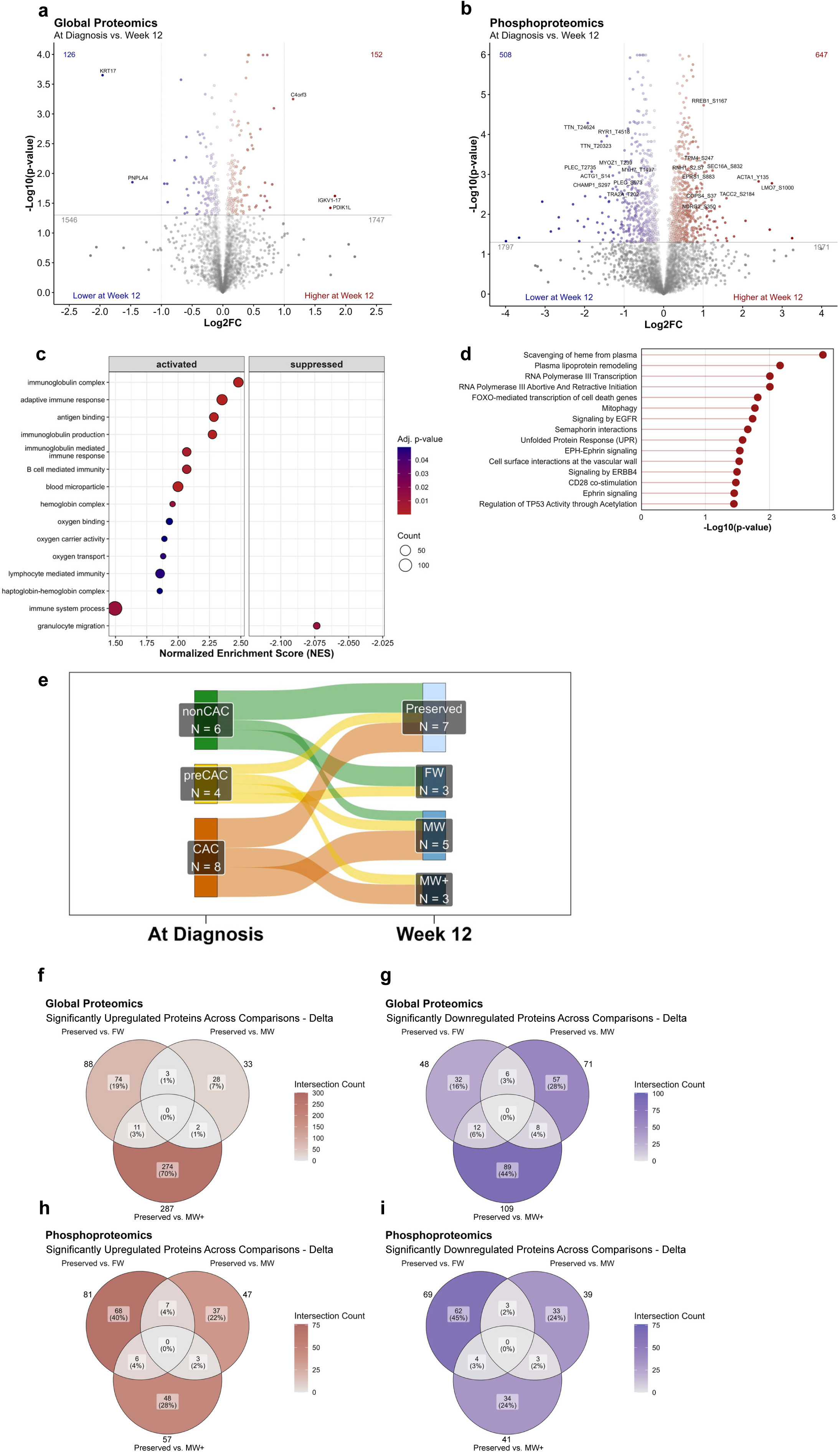
Longitudinal skeletal muscle global and phospho-proteomic profiling during first-line treatment in patients with NSCLC **A:** Volcano plot displaying the log2 fold change (Log2FC) and -log10 p-values of proteins quantified by global proteomics at diagnosis vs. week 12. Significantly regulated proteins with Log2FC > 1 or < −1 are highlighted. Statistical comparisons were performed using paired t-test, and multiple-comparison adjustment was applied using permutation-based correction. **B:** Volcano plot displaying the log2 fold change (Log2FC) and -log10 p-values of proteins quantified by phosphoproteomics at diagnosis vs. week 12. Significantly regulated proteins with Log2FC > 1 or < −1 are highlighted. Statistical comparisons were performed using paired t-test, and multiple-comparison adjustment was applied using permutation-based correction. **C:** Gene set enrichment analysis (GSEA) of log2 fold changes (log2FC) of proteins quantified by global proteomics at diagnosis versus week 12. Analysis was performed using Gene Ontology (GO) categories: Cellular Component (CC), Biological Process (BP), and Molecular Function (MF). The top 15 most enriched pathways are shown. p-values were adjusted for multiple testing using the Benjamini–Hochberg false discovery rate (FDR) method. **D:** Gene-centric Reactome pathway enrichment analysis based on proteins quantified by phosphoproteomics in the comparisons between diagnosis and Week 12. The 15 most significantly enriched pathways are shown. p-values were corrected for multiple testing using the Benjamini–Hochberg (BH) method. **E:** Sankey diagram illustrating group transitions from diagnosis (nonCAC, preCAC, and CAC) to week 12 (preserved, FW = fat wasting, MW = muscle wasting, and MW+ = muscle wasting with additional weight loss). **F**: Venn diagram showing proteins significantly upregulated in the global proteomics across group comparisons at week 12: preserved vs. FW (fat wasting), preserved vs. MW (muscle wasting), and preserved vs. MW+ (muscle wasting with additional weight loss). Statistical comparisons were performed using paired t-test, and multiple-comparison adjustment was applied using permutation-based correction.: Venn diagram showing proteins significantly downregulated in the global proteomics across group comparisons at week 12: preserved vs. FW (fat wasting), preserved vs. MW (muscle wasting), and preserved vs. MW+ (muscle wasting with additional weight loss). Statistical comparisons were performed using paired t-test, and multiple-comparison adjustment was applied using permutation-based correction. **H:** Venn diagram showing proteins significantly upregulated in the phosphoproteome across group comparisons at week 12: preserved vs. FW (fat wasting), preserved vs. MW (muscle wasting), and preserved vs. MW+ (muscle wasting with additional weight loss). Statistical comparisons were performed using paired t-test, and multiple-comparison adjustment was applied using permutation-based correction. **I:** Venn diagram showing proteins significantly downregulated in the phosphoproteome across group comparisons at week 12: preserved vs. FW (fat wasting), preserved vs. MW (muscle wasting), and preserved vs. MW+ (muscle wasting with additional weight loss). Statistical comparisons were performed using paired t-test, and multiple-comparison adjustment was applied using permutation-based correction.

### Proteomic and phosphoproteomic signatures of distinct wasting phenotypes during first-line NSCLC treatment

Having identified broad molecular changes in skeletal muscle, we aimed to identify targetable molecular determinants of muscle-wasting during treatment in NSCLC. Longitudinal paired analysis of 18 patients (6 without CAC, 4 with pre-CAC, and 8 with CAC at diagnosis) across 12 weeks revealed divergent wasting trajectories: throughout NSCLC treatment, 7 patients were weight stable with fat and muscle tissue preserved (*preserved*), 3 patients exhibited fat mass wasting only (*FW*), 5 patients had muscle-wasting only (*MW*), and 3 patients experienced muscle-wasting with weight loss (*MW+*) (Fig 4e). Notably, we observed a clear tendency towards continued muscle-wasting in patients with CAC, while only 1 nonCAC patient exhibited muscle-wasting during first-line NSCLC treatment. In turn, the patients with preCAC showed diverse responses, underscoring the intricacies of cancer cachexia.

Proteomic profiling based on delta values (protein abundance at week 12 minus abundance at diagnosis) revealed distinct treatment-associated remodeling patterns for each patient’s wasting trajectory. For each patient, protein-specific delta changes were compared between patients with wasting and those with preserved muscle mass to identify differential treatment responses. Relative to *preserved* (Fig. 4f,g), patients with *FW* showed significantly divergent regulation of 136 proteins (88 higher, 48 lower), while the *MW* group displayed divergent deltas across 95 proteins (33 higher, 71 lower). In turn, patients in the most severe wasting trajectory (*MW+*) exhibited the greatest proteomic divergence, with 376 proteins significantly altered (287 increased, 89 decreased) relative to *preserved* patients (Fig. 4f,g).

Across the phosphoproteome, delta-based analysis revealed a lessened trajectory-dependent pattern, with 81, 47, and 57 phosphosites showing increased changes in *FW, MW*, and *MW+* patients, respectively (Fig. 4h), and 68, 39, and 41 phosphosites showing decreased changes relative to *preserved* patients (Fig. 4i). Collectively, these between-group delta comparisons reveal a graded intensification of treatment-responsive remodeling across the proteome, whereas changes in the phosphoproteome were comparatively modest. This pattern suggests that progression toward cancer-associated muscle wasting is more tightly linked to persistent alterations in protein abundance than to ongoing phosphorylation dynamics.

### Integrative proteomic and clinical characterization of cancer-associated muscle loss

Having identified distinct proteomic and phosphoproteomic changes associated with different fat and muscle wasting trajectories, we next focused on muscle wasting because of its critical clinical importance for patient outcomes^35,36^. By comparing patients without muscle mass loss (*preserved* and *FW* patients) to those with muscle loss (*MW* and *MW+* patients), we aimed to identify molecular and clinical alterations specifically associated with muscle wasting. Longitudinal analyses of clinical variables (Table 2) demonstrated lower SMI for *MW* and *MW+* patients compared to *preserved* and *FW* patients. During treatment, the nutritional risk score was also lowered, indicating beneficial effects of the treatment (Table 2). Comparing the proteomic delta change from NSCLC diagnosis to week 12, we found that 69 proteins were lower and 68 higher in muscle-wasting patients compared to p*reserved* and *FW* patients (Fig. 5a). GSEA uncovered a distinct proteomic signature, characterized by upregulation of extracellular matrix constituents (Fig. 5d) together with downregulation of mitochondrial proteins (Fig. f) and oxidative metabolism (Fig. 5b-h). Specifically, the most significantly enriched terms within each GO ontology category, revealed Collagen-containing extracellular matrix (GOCC; Fig. 5b), Cell adhesion (GOBP; Fig. 5c), and Extracellular matrix structural constituent (GOMF; Fig. 5d) as the most significantly activated pathways, reflecting pronounced structural and fibrotic remodeling in muscle of NSCLC patients undergoing muscle wasting. In contrast, the most significantly suppressed pathways were Transmembrane transporter complex (GOCC; Fig. 5e), Aerobic respiration (GOBP; Fig. 5f), and NAD(P)H dehydrogenase (quinone) activity (GOMF; Fig. 5g), indicating a coordinated reduction in mitochondrial- and redox capacity–related proteins. These proteomic changes over 12 weeks of first-line NSCLC treatment highlight that patients with muscle wasting undergo distinct molecular adaptations compared with those without wasting. Although we did not stratify by treatment type, the results likely reflect an interplay between tumor- and treatment-driven molecular rewiring, culminating in a fibrotic, energetically impaired muscle phenotype that captures the multifactorial complexity of cancer-associated muscle wasting in humans.

**Figure 5.**
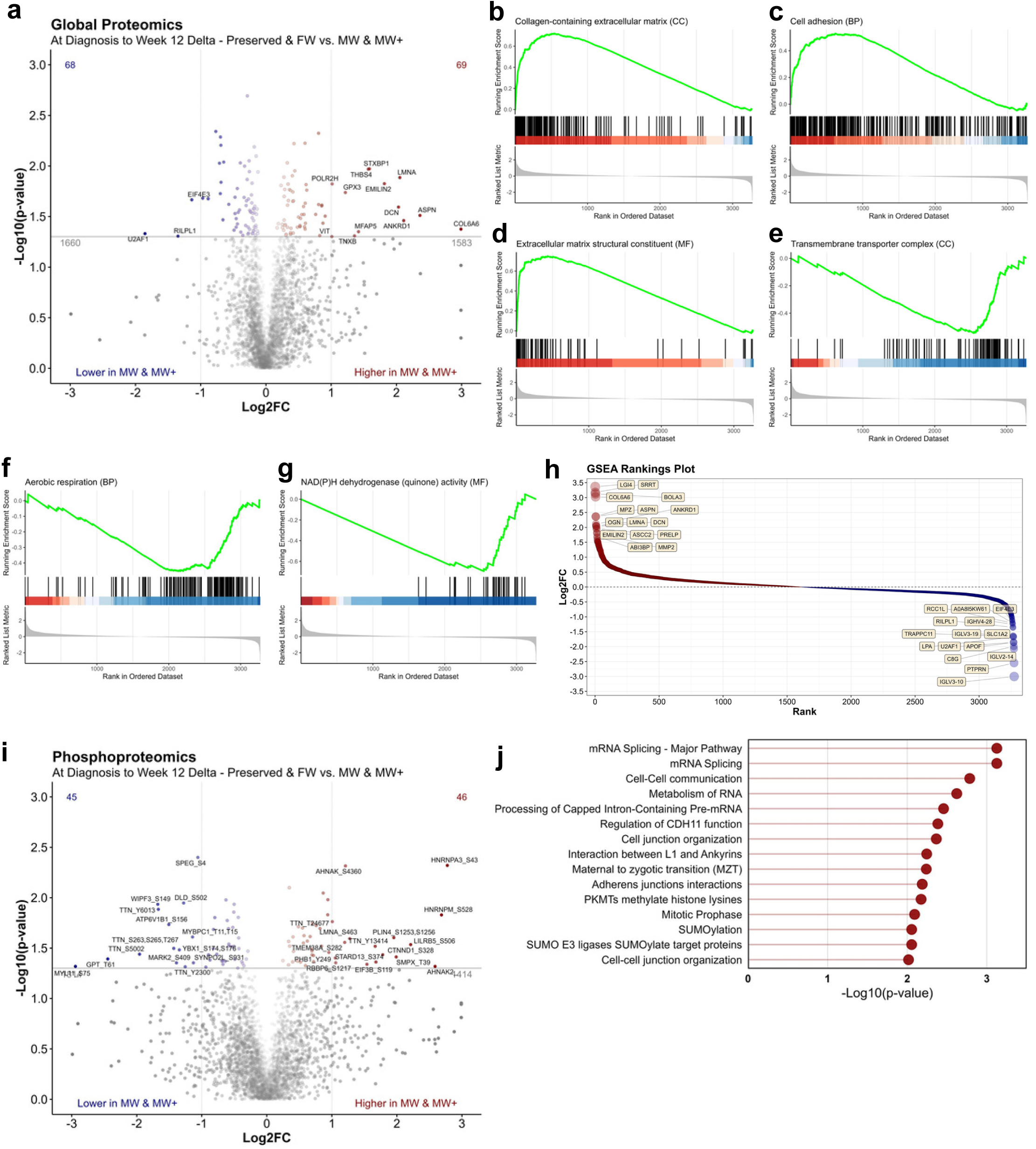
Longitudinal proteomic and phosphoproteomic profiling reveals extracellular matrix and mitochondrial alterations across pooled muscle wasting trajectories in NSCLC **A:** Volcano plot displaying the log2 fold difference (Log2FC) and -log10 p-values of proteins quantified by global proteomics comparing individual delta changes from at diagnosis vs. week 12 between pooled wasting-trajectory groups (preserved/FW vs. MW/MW+). Significantly regulated proteins with Log2FC > 1 or < −1 are highlighted. Statistical comparisons were performed using Welch’s t-test, and multiple-comparison adjustment was applied using permutation-based correction. **B-D:** Gene set enrichment analysis (GSEA) of log2 fold changes (log2FC) of proteins quantified by global proteomics at diagnosis versus week 12. Analysis was performed using Gene Ontology (GO) categories: Cellular Component (CC), Biological Process (BP), and Molecular Function (MF). The most significantly *activated* category within each GO term category was extracted and visualized using GSEA barcode plots. p-values were adjusted for multiple testing using the Benjamini–Hochberg false discovery rate (FDR) method. **E-G:** Gene set enrichment analysis (GSEA) of log2 fold changes (log2FC) of proteins quantified by global proteomics at diagnosis versus week 12. Analysis was performed using Gene Ontology (GO) categories: Cellular Component (CC), Biological Process (BP), and Molecular Function (MF). The most significantly *supressed* category within each GO term category was extracted visualized using GSEA barcode plots. p-values were adjusted for multiple testing using the Benjamini–Hochberg false discovery rate (FDR) method. **H:** Ranking plot showing the Log2FC differences across proteins quantified by global proteomics comparing individual delta changes from at diagnosis vs. week 12 between pooled wasting-trajectory groups (preserved/FW vs. MW/MW+). Top 15 proteins with the highest Log2FC difference are annotated. **I:** Volcano plot displaying the log2 fold difference (Log2FC) and -log10 p-values of proteins quantified by phosphoproteomics comparing individual delta changes from at diagnosis vs. week 12 between pooled wasting-trajectory groups (preserved/FW vs. MW/MW+). Significantly regulated proteins with Log2FC > 1 or < −1 are highlighted. Statistical comparisons were performed using Welch’s t-test, and multiple-comparison adjustment was applied using permutation-based correction. **J:** Gene-centric Reactome pathway enrichment analysis based on proteins quantified by phosphoproteomics comparing individual delta changes from at diagnosis vs. week 12 between pooled wasting-trajectory groups (preserved/FW vs. MW/MW+). The 15 most significantly enriched pathways are shown. p-values were corrected for multiple testing using the Benjamini–Hochberg (BH) method.

**Table 2.** Clinical, functional, and body composition variables across pooled wasting trajectories in week 12 Comparison of changes in clinical variables between preserved & fat wasting (FW) patient vs. muscle wasting (MW) & muscle wasting with additional weight loss (MW+) patients across 12 weeks. Data are presented as n, mean (SD = standard deviation) and p value between groups. Statistical comparisons were performed using Welch’s t-test. **Abbreviations:** CRP, C-reactive protein; BMI, body mass index; HgbA1c, glycated hemoglobin; AU, arbitrary units.

|  | Preserved/FW | MW/MW+ | P value |
| --- | --- | --- | --- |
| **Sit to stand test** (cycles/30 sec) | N = 9 | N = 8 | 0.32 |
| Mean (SD) | 1.78 (3.03) | 0.25 (3.11) |  |
| **Fitness level** (mL O<sub>2</sub>/min/kg) | N = 7 | N = 4 | 0.36 |
| Mean (SD) | -2.57 (6.08) | 1.25 (6.18) |  |
| **Hand Grip Strength** (kg) | N = 9 | N = 8 | 0.69 |
| Mean (SD) | -0.28 (0.93) | -1.25 (6.20) |  |
| **BMI** (kg/m<sup>2</sup>) | N = 10 | N = 8 | 0.13 |
| Mean (SD) | 0.89 (0.93) | -1.02 (3.06) |  |
| **Adipose Tissue Mass Index** (kg/m<sup>2</sup>) | N = 9 | N = 8 | 0.07 |
| Mean (SD) | 0.49 (0.88) | -0.92 (1.81) |  |
| **Lean body Mass Index** (kg/m<sup>2</sup>) | N = 9 | N = 8 | 0.84 |
| Mean (SD) | 0.07 (0.98) | -0.05 (1.31) |  |
| **Skeletal Muscle Radiation Attenuation** (HU) | N = 9 | N = 8 | 0.74 |
| Mean (SD) | -0.36 (7.17) | 0.68 (5.20) |  |
| **Skeletal Muscle Index** (cm<sup>2</sup>/m<sup>2</sup>) | N = 9 | N = 8 | <0.001 |
| Mean (SD) | 2.14 (1.60) | -6.76 (4.43) |  |
| **Appendicular Skeletal Muscle Index** (kg/m<sup>2</sup>) | N = 9 | N = 8 | 0.53 |
| Mean (SD) | 0.15 (0.43) | -0.09 (0.91) |  |
| **Subcutaneous Adipose Tissue Index** (cm<sup>2</sup>/m<sup>2</sup>) | N = 9 | N = 8 | 0.15 |
| Mean (SD) | 5.09 (15.06) | -4.93 (12.31) |  |
| **Nutritional Risk Score** (AU) | N = 10 | N = 8 | <0.01 |
| Mean (SD) | 0.90 (15.06) | -4.25 (3.96) |  |
| **Hemoglobin** (mmol/L) | N = 10 | N = 8 | 0.88 |
| Mean (SD) | -0.66 (1.45) | -0.76 (1.31) |  |
| **HbA1c** (mmol/mol) | N = 8 | N = 4 | 0.54 |
| Mean (SD) | -0.25(6.39) | 2.00(5.35) |  |
| **Albumin** (g/L) | N = 10 | N = 8 | 0.78 |
| Mean (SD) | 1.60 (5.97) | 0.88(4.88) |  |
| **CRP** (mg/L) | N = 9 | N = 8 | 0.2 |
| Mean (SD) | -16.11 (22.18) | 2.03 (31.48) |  |

Interestingly, while the global proteome revealed increased extracellular matrix remodeling and reduced oxidative capacity, phosphoproteomic analysis highlighted post-translational phosphorylation changes in pathways governing RNA processing, cell junction organization, and chromatin regulation. Comparing the delta change in the muscle phosphoproteome from NSCLC diagnosis to week 12, we found that 46 phosphoproteins were upregulated and 45 downregulated in the muscle-wasting patients compared to *preserved* and *FW* patients (Fig. 5i). The top Reactome pathways (Fig. 5j) include mRNA splicing, cell–cell communication, metabolism of RNA, and cell junction organization, suggesting extensive modulation of transcriptional and structural signaling networks in muscle wasting. These phosphorylation-driven changes imply that, in addition to metabolic and fibrotic remodeling, muscle wasting during NSCLC treatment is associated with broad post-translational reprogramming that affects gene expression and cytoskeletal integrity. Together, the proteomic and phosphoproteomic analyses reveal a multilayered adaptive response in which tumor- and treatment-induced stress remodel both structural and regulatory components of skeletal muscle, culminating in a fibrotic and metabolically compromised molecular phenotype specific to patients with muscle wasting.

### Common and distinct molecular CAC responses in muscle of female and male patients with NSCLC

After identifying proteins and phosphosites linked to CAC at diagnosis and muscle loss during treatment, we investigated sex-specific molecular signatures in NSCLC patients, motivated by recent RNA-sequencing studies showing that males and females share only ∼50% of their transcriptomic alterations^18^. Although the non-CAC and CAC groups were sex-matched, stratifying for sex could reveal shared and sex-specific adaptations in skeletal muscle, a largely unexplored area in human cachexia and muscle wasting.

In the global proteomics dataset, stratification by sex revealed 63 proteins upregulated and 106 downregulated in CAC males compared with nonCAC males (Fig. S4a), whereas 122 proteins were elevated and 21 reduced in CAC females compared with nonCAC females (Fig. S4b) at diagnosis. In the phosphoproteome, using the same stratification, 100 phosphosites were increased and 72 decreased in CAC males (Fig. S4c, while 94 phosphosites were increased and 79 decreased in CAC females (Fig. S4d). Notably, there was almost no overlap between male and female responses to CAC in either the global proteome or the phosphoproteome, illuminating highly sex-specific molecular muscle adaptations to NSCLC in humans (Fig. S4e-h). Only one protein, EMC2 (ER membrane protein complex subunit 2), a core subunit of the endoplasmic reticulum membrane complex, which is involved in membrane protein insertion and ER homeostasis^37^, was upregulated in both sexes with CAC compared with non-CAC patients (Fig. S4e). Integrating proteomic, clinical, and functional data, these findings define CAC- and muscle-specific molecular signatures marked by sex-dependent remodeling and perturbation of calcium signaling pathways.

## DISCUSSION

By integrating global and phosphoproteomic changes in skeletal muscle with comprehensive clinical data on physical activity, body composition, muscle quality, plasma biomarkers, and patient-reported nutritional risk, we identified key muscle proteins and phosphorylation sites associated with CAC at diagnosis, as well as with muscle wasting during treatment. These findings identify novel molecular targets in muscle, advancing our understanding of CAC that can guide therapeutic strategies to combat muscle wasting.

This integrated analysis yielded five major insights. First, we identified distinct proteomic and phosphoproteomic signatures involving calcium, anabolic, and stress signalling, alongside extracellular matrix and mitochondrial alterations that differentiate patients with and without CAC at the time of NSCLC diagnosis. Second, we found that first-line treatment markedly upregulated inflammatory pathways, indicating a pronounced treatment effect on the muscle proteome and phosphoproteome. Third, we delineated molecularly distinct trajectories of muscle wasting throughout treatment, which led to the fourth key finding, that patients exhibiting muscle loss showed specific regulation of proteins involved in extracellular matrix composition, mitochondrial function, and mRNA-related processes. These findings complement our parallel cross-sectional study of NSCLC skeletal muscle microenvironment biology, which revealed extracellular matrix remodeling, mitochondrial alterations, calcium handling disruptions, immune and inflammatory alterations, and shifts in FAP populations at diagnosis^38^. Finally, exploratory analysis of possible sexual dimorphism suggested notable sex-specific molecular responses linked to CAC and muscle wasting, pointing to a previously underappreciated layer of biological regulation..

Our first finding provides the first comprehensive characterization of proteomic and phosphoproteomic signatures distinguishing NSCLC patients with and without CAC at diagnosis, generating the first open-source muscle *CACProteome&Phosphoproteome* dataset. The limited prior human work^39–41^ revealed structural and metabolic protein alterations in patients, yet lack detailed phosphoproteomic profiling, matched CAC vs nonCAC comparisons, and integration with clinical measures and muscle function parameters. By combining global and phosphoproteomic analyses with multiple clinical variables, including physical activity, body composition, muscle quality, plasma biomarkers, and nutritional risk, our study addresses these gaps, defining molecular features of CAC directly in patients and across treatment.

Our global proteome analysis revealed clinically relevant CAC-associated changes mainly in proteins regulating myocellular calcium handling at NSCLC diagnosis. Guided by an enrichment analysis and GO term associations from a concept network^42^ we identified highly CAC regulated proteins related to calcium handling that highly correlated with clinical variables at diagnosis suggesting that altered calcium handling translates into clinical outcomes affecting the patients directly. This aligns with a recent proteomic study in pancreatic cancer, which, despite lacking a non-CAC comparison, also reported pronounced perturbations in endoplasmic reticulum components in rectus abdominis muscle^43^. With the identification of marked calcium-related perturbations, which was also recently suggested in preclinical CAC^44^, these findings corroborate a previous study reporting alterations in muscle contractile myosins and mitochondrial proteins in five CAC patients compared with weight-stable patients with mixed cancers^39^. Another study revealed transcriptional alterations in genes involved in protein synthesis, neuronal function, extracellular matrix, immune-related inflammation, and metabolism in pancreatic and colorectal CAC^18^. Building on our proteomic analyses, further deep phosphoproteomic profiling uncovered pronounced perturbations in posttranslational regulation of proteins related to energy- and growth, alongside disrupted insulin signaling, consistent with impaired anabolic control and ongoing muscle loss in CAC. The regulated phospho-sites also correlated with key clinical variables, supporting the premise of disrupted proteomic and phosphoproteomic regulation as core in CAC at diagnosis. This comprehensive mapping of the human muscle proteome, including post-translational modifications and correlations with key clinical variables at diagnosis, is unique and provides a foundation for identifying therapeutic targets in CAC. Collectively, our findings reveal a complex, clinically relevant interplay between calcium signaling and muscle growth pathways, offering novel and clinically anchored mechanistic insights into the molecular drivers of CAC.

Enabled by repeated muscle biopsies obtained at week one and week 12 during first-line therapy, our second key finding, revealed pronounced treatment-induced remodeling of the skeletal muscle, particularly proteins involved in inflammatory pathways. Phosphoproteomic analyses further illuminated posttranslational regulation of pathways related to mitophagy, FOXO-mediated transcription of cell death, and RNA transcriptional regulation, indicating disruption of transcriptional control. Our results corroborate data from seven patients with head and neck or cervical cancer following seven weeks of chemotherapy, where the predominant effects involved metabolic processes, cytoskeletal organization, oxygen transport, and apoptotic signaling^45^. Given that most patients in our cohort received immune checkpoint inhibitors, our data suggest that this treatment exerts a profound, previously underappreciated impact on proteins involved in skeletal muscle inflammatory signaling and transcriptional regulation.

By leveraging the longitudinal design of our study, we delineated distinct trajectories of fat and muscle wasting during NSCLC treatment. These analyses revealed a consistent pattern of progressive muscle loss in 62.5 % of patients with CAC, whereas none of the non-CAC patients exhibited body weight loss with muscle wasting. Although not evaluated in our study, recent work in pancreatic cancer has identified distinct CAC phenotypes that independently associate with survival outcomes^46^. Although that study lacked longitudinal body composition data and molecular profiling, our findings suggest, together with theirs^46^, that distinct molecular mechanisms underlie each wasting trajectory, with potential implications for survival.

By coupling individual wasting trajectories with molecular profiling, our third finding uncovered distinct, treatment-associated muscle remodeling patterns for each trajectory, most prominently reflected at the proteomic level. This finding suggests that progression toward cancer-associated muscle wasting is more tightly linked to alterations in protein abundance than to dynamic changes in phosphorylation. Importantly, each patient’s molecular trajectory was referenced to their baseline molecular signature, at which time CAC and non-CAC patients already displayed pronounced differences. As most CAC patients continued to lose muscle mass while non-CAC patients remained weight stable, our data could suggest that early proteomic and phosphoproteomic alterations in calcium and muscle growth pathways may have predisposed the muscle to further proteomic reprogramming and wasting upon initiation of systemic therapy.

Our fourth major finding was the identification of molecular proteomic and phosphoproteomic changes specific to the skeletal muscle wasting trajectories throughout treatment. Here, proteomics revealed regulation of proteins related to extracellular matrix constituent as well as NADPH dehydrogenase activity, while the phosphoproteome revealed enrichment of mRNA related pathways and cell communication, which jointly may suggest an interaction between the ECM, mitochondrial and muscle wasting mechanics. As all but one patient on the muscle-wasting trajectories were CAC or pre-CAC at diagnosis, we speculate that the baseline proteomic and phosphoproteomic alterations in calcium and muscle growth pathways predispose skeletal muscle to progressive wasting during treatment. Thus, beyond revealing the molecular rewiring underlying muscle loss, these findings underscore the high risk of further muscle wasting in CAC patients and highlight potential pathways for early intervention.

Finally, our results indicate striking sex-specific proteomic and phosphoproteomic responses associated with CAC at diagnosis and muscle wasting through treatment. Sexual dimorphism was also recently noted at the transcriptomic level in rectus abdominis muscle of pancreatic and colon cancer CAC patients^18^. Notably, at NSCLC diagnosis, only a single protein, the endoplasmic reticulum membrane complex protein, EMC2, was associated with CAC in both men and women. Given that the endoplasmic reticulum in skeletal muscle is contiguous with the sarcoplasmic reticulum, these alterations may disrupt calcium handling, linking endoplasmic reticulum stress to impaired calcium signaling and muscle wasting in CAC across sexes. Interestingly, male cancer patients, incl. NSCLC^47^ generally have higher prevalence of cachexia, greater weight loss or muscle wasting, and worse outcomes compared with females^48^. Our study underscores the need for further molecular investigations of sex-specific differences in CAC. Collectively, these findings reveal sexual dimorphism as an underappreciated layer of biological regulation in cachexia, which must be addressed to advance personalized therapeutic strategies.

Key strengths and limitations should be considered when interpreting these data. This study is strengthened by the integration of global and phosphoproteomic profiling with deep clinical phenotyping, including CT-derived body composition, anchoring molecular insights in clinically relevant patient measures. Our longitudinal design, enabled by repeated muscle biopsies and comprehensive functional assessments in newly diagnosed advanced-stage NSCLC patients, minimized heterogeneity and allowed characterization of individual muscle-wasting trajectories. Uniform tissue sampling and rigorous clinical data collection at a single center further ensured high-quality data. Limitations include the modest cohort size and focus on a single cancer type, which constrain generalizability and preclude definitive conclusions on sex-specific differences or cross-cancer comparisons, leaving open whether CAC is molecularly conserved or cancer-specific. Functional validation of identified targets was not performed. Additionally, skeletal muscle heterogeneity, including multinucleated fibers, immune and endothelial cells, stem cells, and mesenchymal progenitors, limits resolution, emphasizing the need for spatial or single-cell proteomic approaches to capture cell type–specific alterations.

All together, our integrated analysis of repeated muscle biopsies from patients with advanced NSCLC identifies disrupted calcium-handling signaling as a central feature of CAC at diagnosis that correlated with multiple key clinical patient measures. By linking longitudinal proteomic and phosphoproteomic profiles with clinical measures, including body composition, muscle quality, physical activity, plasma biomarkers, and nutritional risk, we provide molecular clinically anchored insights to the pathways that underpin progressive muscle loss. Our findings highlight calcium homeostasis as a potential therapeutic target to prevent or mitigate muscle wasting in NSCLC.

## METHODS

### Subjects

Patients were included between August 2022 to October 2023 from Department of Oncology, Copenhagen University Hospital – Rigshospitalet within the first week after being diagnosed with advanced stage NSCLC;. Inclusion criteria were: (1) age > 18 years, (2) pathologically confirmed NSCLC, TNM v. 8.0 stage IIIb/IV not eligible to concurrent chemo/radiation therapy as primary treatment, (3) referred for first-line palliative systemic anticancer therapy (4) having a staging/baseline CT within 4-6 weeks of initiation of treatment, or a baseline scan planned within the first week of treatment, (5) being in Eastern Cooperative Oncology Group Performance Score (ECOG PS) 0-2, (5) having signed the informed consent form to the study. Exclusion criteria were: (1) any other known malignancy requiring active treatment, (2) palliative radiotherapy as primary treatment, (3) ECOG PS >2, (4) physical disabilities excluding physical testing, (5) inability to understand Danish, (6) inability to understand scoring systems/patient-reported outcome measures, (7) taking anti-coagulant therapy that could not be discontinued for the muscle biopsy. Patients eligible for anticancer treatment were not excluded due to other comorbidities. Patients were classified, according to self-reported weight loss over 6 months prior to their diagnoses as: CAC (more than 5% loss of stable body weight over the past 6 months, or a BMI less than 20 kg/m² and ongoing weight loss of more than 2%, or sarcopenia and ongoing weight loss of more than 2% but have not entered the refractory stage^2^; pre-CAC (1-5% WL), or nonCAC (increase in weight or up to 1% WL). Sarcopenia was defined as L3 CT-derived skeletal muscle index (SMI) (cm2/m2) < 43/41 for normal or underweight men/women and <53/41 for overweight and obese men/women) ^2,5^. In one patient we could not obtain CT-derived SMI measures why DXA-derived lean and fat mass change was used instead. In the global and phospho-proteomic analyses at diagnosis, only patients having CAC and nonCAC were included in the analysis, given the low n (=4) of patients in the pre-CAC group.

### Protocol, approvals and handling of data

The study was carried out in accordance with the Helsinki Declaration, approved by the Regional Scientific Ethical Committee of the Capital Region of Denmark (H-21035808), and registered at www.clinicaltrials.gov (NCT05307367). All clinical data has been stored in a Redcap server operated by the Capital Region of Denmark and in agreement with the European Union’s General Data Protection Regulation (GDPR). For this a data transfer agreement was obtained (Jr.nr 22019423). Approval of processing of data was given from the local institutional board (<u>514-0685/22-3000).</u> Witten informed consent was obtained from each patient before participation.

### Physical functioning, body composition and physical activity

In week 1 and week 12 patients were invited to the MuscleLab at Copenhagen University Hospital, Rigshospitalet for analyses of body anthropometry and composition with DXA-scan. Besides this, functional testing of maximal hand grip strength, sit-to-stand test 30s^49^ and VO2 peak were recorded^50,51^. From both the diagnostic CT scan and the evaluating CT-scan body composition was calculated for the patients. Skeletal muscle index (SMI) and subcutaneous adipose tissue (SATI) were extracted by normalizing the tissue area on L3 level to patient height (cm^2^/meter^2^). Mean skeletal muscle radio attenuation (SMRA) of the skeletal muscle area was calculated and used as a marker for muscle quality/myosteatosis^52^. Based on the specific loss of different body compartment index (SMI and SATI) as well as body weight loss between the diagnostic CT scan and the evaluating CT scan in week 12, patients were grouped into specific wasting phenotypes defined as follows; ***preserved*** (no wasting) or <2.5% loss of SMI and SATI, ***FW*** (fat-wasting): loss of SATI >2.5%, ***MW*** (muscle-wasting): loss of SMI >2.5 %, and ***MW+*** (muscle-wasting + weight loss): loss of SMI >2.5 % and weight loss >2.5%.

Physical activity was conducted with the SENS motion activity measurement system (SENS motion system)^53^; that measures movement continuously at 25 Hz (every 2,5 s), 24 h a day. The accelerometer was placed on the lateral side of the right thigh with a SENS patch (art.no 136-100-B). Accelerometer data from week 1 was used. A predefined algorithm integrates the sensor’s orientation concerning gravity and the recorded acceleration and categorizes the recordings into predefined types of physical activities: ‘resting’ (lying down, sitting) and ‘time-on-feet’ (standing, sporadic/continuous walking, running, cycling). The intensity counts and time-on-feet from 6 am. to 10 pm. was calculated. Validated in other sedentary patient groups, it has shown good and consistent ability to distinguish between activity, standing, and sedentary behavior^54,55^.

### Patient-reported outcome measures

An online platform retrieving patient-reported outcome measures (PROMs) was linked to the Redcap server. All patients were reporting nutritional risk scores every third week to ensure individualized nutritional and supportive care, and to minimize secondary nutritional impact symptoms for the patients. In this study we specifically used the patient-reported outcome measure PG-SGA scores in week 1 and week 12^56,57^.

### Blood sample and muscle tissue collection

The day before the first systemic treatment was initiated in week 1, baseline blood samples were taken non-fasting in the Department of Oncology, Copenhagen University Hospital – Rigshospitalet. This again 12 weeks into first-line treatment. Analyses were performed at the Department of Clinical Biochemistry, Copenhagen University Hospital – Rigshospitalet; and from here hemoglobin, HgbA1c, albumin and CRP are used in this analysis^58^. In the same first week of systemic treatment the patients were invited after an overnight fast to the first muscle biopsy; obtained median day 6 after initiating first-line treatment. The second biopsy was obtained median Day 89 (Table 1). Blood sampling, nutritional risk assessment, physical function testing, and body composition analysis were conducted at the same timepoints.

Patients were urged not to do exercise 48 hrs. prior to the biopsies. Percutaneous biopsy of the left vastus lateralis was performed at 2/3 from the most proximal aspect of the patella and the inguinal fold. Skin was sterilized (2% chlorhexidine-70% isopropyl alcohol) and the site was infiltrated with local anesthetic (lidocaine without adrenalin 20mg/ml). The procedure from here on was performed sterile with chlorhexidine 5% and surgical covering. A 6-mm incision was made through the skin, fat, and down to fascia, and the biopsy needle (6 mm Bergström) was advanced past the fascia into the muscle. Suction was applied and the tissue was obtained. All biopsies were rinsed in ice cold saline within 30 s after procedure, and within 1-2 min sufficient tissue was snap frozen in LN2. Follow up biopsies were conducted in week 12 (median day 89) in all 18 patients. Week 12 biopsies were conducted on the right leg.

### Immunohistochemistry

Muscle biopsies were prepared as described in Battey et al. 2025^38^. In short, muscle biopsies were embedded in optimal cutting temperature (OCT) compound and cryosectioned at 8 μm thickness with fibers oriented perpendicular to the blade to ensure accurate assessment of muscle fiber morphology and size. Sections were blocked in 5% donkey serum and incubated with primary antibodies for 3 hours at room temperature. Corresponding secondary antibodies were used: DyLight 488–conjugated anti–mouse IgG1 (Jackson #115-545-205), DyLight 405–conjugated anti–mouse IgM (Jackson #715-475-020), and Alexa Fluor 568–conjugated anti–rabbit (Invitrogen #A11011). For histology, antibodies recognizing Myosin Heavy Chain 2 (1:100 A.74, DSHB), Myosin Heacy Chain 7 (1:100 A4.840, DSHB) and Laminin (1:200, PA1-16730, Invitrogen) were used.

### Sample preparation for proteomic analysis

After storage at −80°C a fraction of each skeletal muscle biopsies in LN2 was crushed and 5-7 mg was transferred to Twintec plates for global and phospho-proteomic analyses. Tissue samples were lysed by addition of preheated (95 °C) lysis buffer (5% SDS, 100 mM Tris pH 8.5) and lysed with a BeatBox homogenizer for 5 min at high setting. Samples were incubated at 95 °C for 10 min and lysed with a second round of BeatBox for 10 min at high setting. Samples were reduced using 5 mM TCEP for 15 min at 55 °C and alkylated with 20 mM CAA for 30 min at room temperature. Next, 233 μL acetonitrile (ACN) and 25 μL of pre-equilibrated (70% ACN) MagReSyn Hydroxyl beads (Resyn Biosciences) were added to 100 μL protein extract (140 μg protein). Samples were mixed and settled for 10 min at room temperature. Beads were washed three times with 1 mL 95% ACN, twice with 1 mL 70% EtOH, and digested over night with 150 μL digestion buffer (50 mM TEAB) containing Trypsin and Lys-C in a 1:50 enzyme to protein ratio. Samples were acidified with 50 μL 10 % formic acid, and 200 ng loaded onto EvoTips for global proteome quantification.

Phosphopeptide enrichment was carried out on a KingFisher Apex (Thermo) instrument in 96 well format as described^59^. Acidified digested peptides were vacuum centrifuged to near dryness, resuspended in 200 μL loading buffer (0.1 M glycolic acid, 80% ACN, 5% TFA), and incubated with 20 μL of washed and equilibrated 1:1 mix of Ti-IMAC and Zr-IMAC beads (20 mg/ml, Resyn Biosciences) for 20 min at medium speed. Beads were washed once with 500 μL loading buffer, once with 500 μL wash buffer 1 (80% ACN, 5% TFA), and once with 500 μL wash buffer 2 (10% ACN, 0.2% TFA), for 2 min at fast speed. Phosphopeptides were eluted with 100 μL 1% ammonia for 10 min at medium speed, and the eluate acidified by addition of 20 μL 10% formic acid. Peptides were vacuum centrifuged to near dryness, resuspended in 1% TFA, and loaded onto EvoTips.

### Data acquisition by liquid chromatography–mass spectrometry LC-MS

Peptides were separated on a Pepsep 15 cm, 150 μM ID column packed with C18 beads (1.5 μm) using an Evosep ONE HPLC system applying the default 30-SPD (30 samples per day) method. The column temperature was maintained at 50 °C. Upon elusion, peptides were injected via a Nanospray Flex ion source and 30-uM steel emitter (Evosep) into a Tribrid Ascend mass spectrometer (Thermo Scientific). Spray Voltage was set to 2200 (V). Data was acquired in data independent mode with the Orbitrap MS resolution 120.000 for full scan range 450-1150 m/z, AGC target was 250% and maximum injection time 45 ms. A total of 49 DIA scans with 13.7 Th width and 1 Th overlap spanning a mass range of 472-1143 m/z were acquired at 15,000 MS resolution, AGC target 1000% and maximum injection time of 27 ms. HCD fragmentation normalized collision energy (NCE) was set to 27%.

### Protein identification by computational data analysis

MS files were processed using Spectronaut version 18 (Biognosys) in direct DIA search mode with the default “BGS Phospho PTM Workflow.” The homo sapiens FASTA database was used. Carbaminomethylation of cysteine was set as a fixed modification, while oxidation of methionine, acetylation of the protein N-terminus, and phosphorylation of tyrosine, serine, and threonine were set as variable modifications. The maximum number of missed cleavages was set to 2, and the minimum peptide amino acid length was set to 7. PTM localization was activated, and the site confidence score cutoff for phosphoproteomics data was set to 0.75. The false discovery rate for PSM, peptide, and protein groups was set to 0.01.

### Bioinformatic processing of mass spectrometry data

All analyses were performed using RStudio (v4.3.1). A threshold of n ≥ 3 per group was applied for group-wise comparisons. Data were log2-transformed, and missing values were not imputed; therefore, analyses requiring complete datasets (e.g., principal component analysis) were conducted using only proteins without missing values. For unpaired comparisons, two-sided Welch’s t-tests were used to compute p-values, while a paired t-test was used in the paired analyses. To assess the empirical p-value distribution, up to 15,000 permutations were conducted per test. The permuted p-value was calculated by dividing the number of permutation-based tests with a p-value lower than the observed p-value by the total number of permutations. If the permuted p-value exceeded the original p-value it replaced the original value. No adjustment for multiple comparisons was applied to the sex stratified analyses (male nonCAC vs. male CAC and female nonCAC vs. female CAC) due to the exploratory nature of the comparisons. Gene Set Enrichment Analysis (GSEA) and Over-representation analysis (ORA) were performed using the R package *clusterProfiler* using Homo sapiens as reference organism, with all quantified proteins used as background. Gene Ontology enrichment was conducted across three categories: cellular component (GOCC), biological process (GOBP), and molecular function (GOMF). Enrichment was assessed using a hypergeometric test with Benjamini-Hochberg correction for multiple comparisons. Redundant GO terms were collapsed using a similarity threshold of 0.7. Gene-centric phosphoproteome enrichment was performed as rank-based pathway enrichment using the R package *PhosR* against curated Reactome pathways. Kinase perturbation analysis was performed using the R package *directPA*, against known kinase-substrate relationships to test whether phosphorylation changes of a kinase substrates were directionally biased across conditions. Kendall’s tau rank correlation was used for all correlation analyses.

### Statistics

Data are expressed as mean ± SE, including individual values where applicable. Significance was set at α=0.05. Log2FC comparisons between groups were made using a paired student’s T-test or Welch’s test applying permutation-based correction to account for multiple hypothesis testing. Data plots were generated using R Studio (4.5.0) and GraphPad Prism 10.1.1.

## Data and Code Availability Statement

The mass spectrometry proteomics data will be deposited to the Perseus bioinformatics Platform for Integrative Analysis of Proteomics Data^60^. All data are available upon reasonable request.

## ACKNOWLEDGEMENT

We thank research assistant Nanna Hahn for her dedicated work throughout the study with data collection and direct patient contact and we thank both Nanna Hahn and research assistant Nicoline Resen Andersen for an invaluable effort in the handling of biological samples. We thank nurse Marie Aller and Fie Sylvest and physiotherapist Magnus Nygaard Bech for dedicated work with patient flow and patient support as well as physiotherapist Malthe Mollerup-Degn Hermann, Andreas Ubberup Larsen and Barbara Bordignon for the dedicated work and support for the patients during the physical assessments. We thank all nurses and medical doctors at the Department of Oncology, Copenhagen University Hospital, Rigshospitalet for contributing to successful inclusion and support to patients and relatives who participated in the study. We thank senior radiology consultant Thomas Skårup Kristensen for crucial guidance in acquisition of radiology data. We thank assistant professor Wouter van de Worp for support and guidance regarding handling and interpretation data, as well as research assistant Ralph Brecheisen for body composition analysis. Proteomics sample preparation, LC-MS acquisition, and proteomic data analysis were carried out by the Proteomics Research Infrastructure (PRI) at the University of Copenhagen (UCPH), supported by the Novo Nordisk Foundation (NNF) (grant agreement number NNF19SA0059305). We thank collaborating colleagues from PRI Michael Wierer, Paola Pisano and David Oliver Schlessinger. Finally, and foremost, we express our deepest gratitude to all participating patients and relatives making this research study possible.

## AUTHORS CONTRIBUTIONS

Conceptualization: J.S. and L.S. Data curation: J.S., E.B., A.I., Z.O., C.V., N.Ø. and L.S. Formal analysis: J.S., C.V., Z.O., J.M., E.B., A.I., O.D., N.Ø. and L.S. Funding acquisition: J.S. and L.S. Investigation: J.S. Methodology: J.S., E.B., O.D., and L.S. Project administration: J.S. and L.S. Resources: J.S. and L.S. Software: J.S., C.V., O.D., N.Ø., and L.S. Supervision: L.S. E.A.R. Validation: J.S. and O.D. Visualization: J.S., C.V., E.B., Z.O., and O.D. Writing original draft: J.S., C.V., Z.O., and L.S. Writing review & editing: all authors.

---

**Figure S1.**
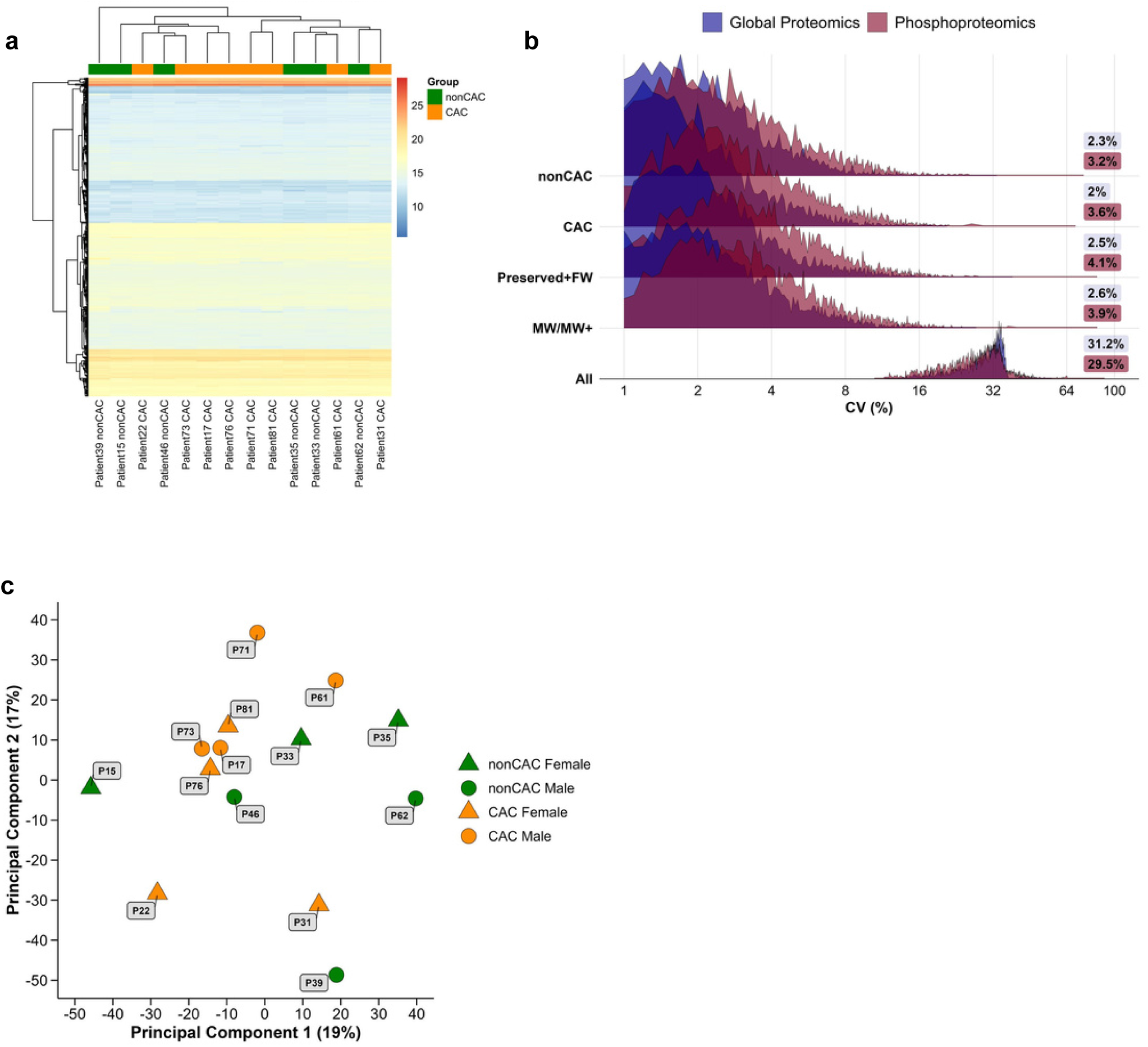
Global proteomics; CAC vs. nonCAC at diagnosis in patients with NSCLC and CV% across analyses. **A:** Hierarchical clustering of protein quantified in the global proteomics analysis based on log2-transformed intensities in nonCAC and CAC patients at diagnosis. **B:** Distribution of coefficients of variation (CV%) across the global- and phosphoproteome in nonCAC, CAC, preserved/FW and MW/MW+ patients. The x-axis has been log2-transformed to increase visibility and numbers in coloured boxes indicate median CV%. **C:** Principal component analysis of protein quantified in the global proteomics analysis showing sample clustering across biological sex and nonCAC and CAC.

**Figure S2.**
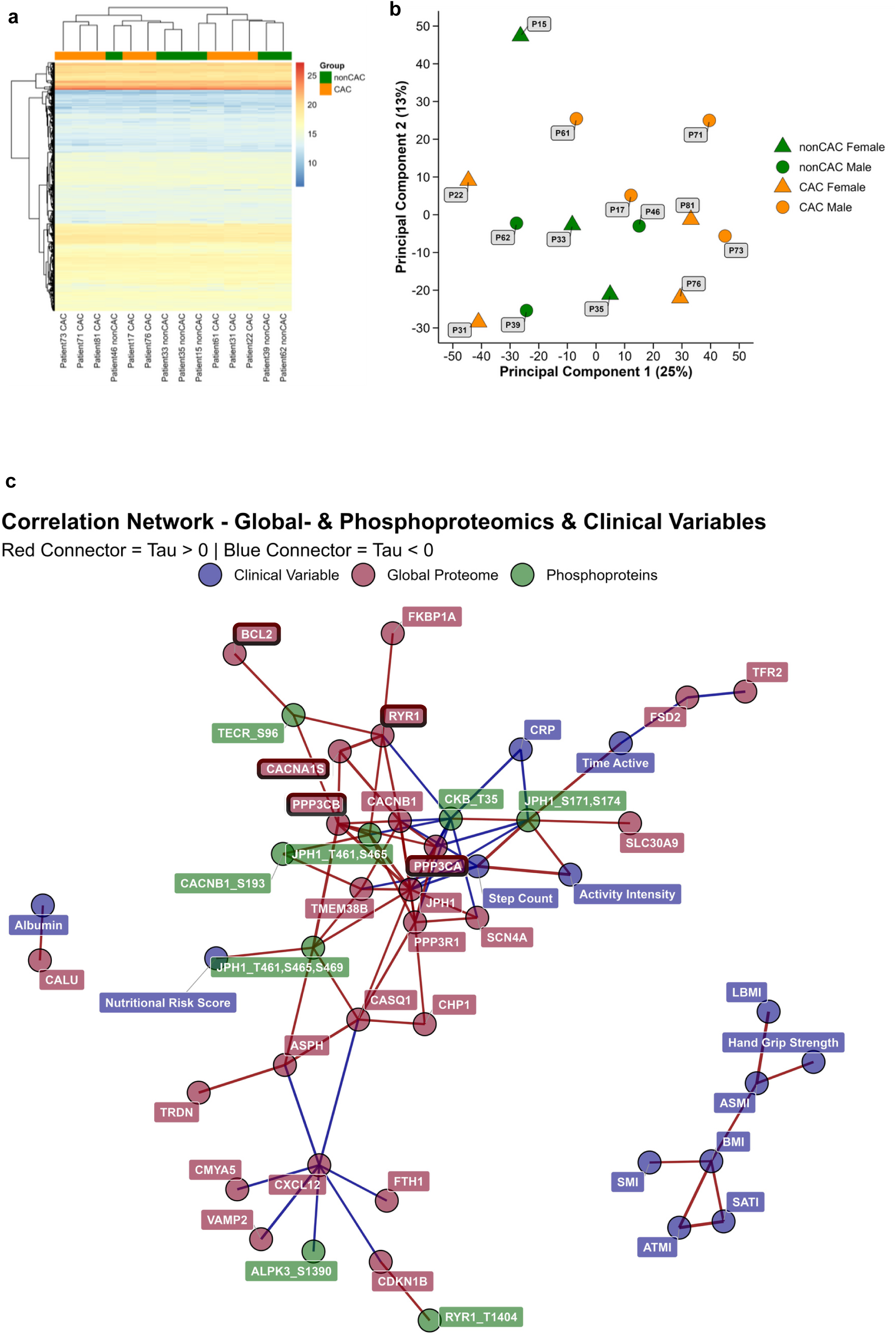
Phosphoproteomics; CAC vs. nonCAC at diagnosis in patients with NSCLC and correlation network analysis. **A:** Hierarchical clustering of protein quantified in the phosphoproteomics analysis based on log2-transformed intensities in nonCAC and CAC patients at diagnosis. **B:** Principal component analysis of protein quantified in the phosphoproteomics analysis showing sample clustering across biological sex and nonCAC and CAC. **C:** Correlation network analysis integrating four feature categories: (1) clinical variables, (2) phosphosites significantly altered between nonCAC and CAC patients at diagnosis whose corresponding proteins were also significantly regulated in the global proteome, and (3) proteins identified within the concept network (Fig. 2c). Only nodes exhibiting Kendall’s correlation coefficient > 0.6 or < −0.6 with at least one other feature were included in the network visualization.

**Figure S3.**
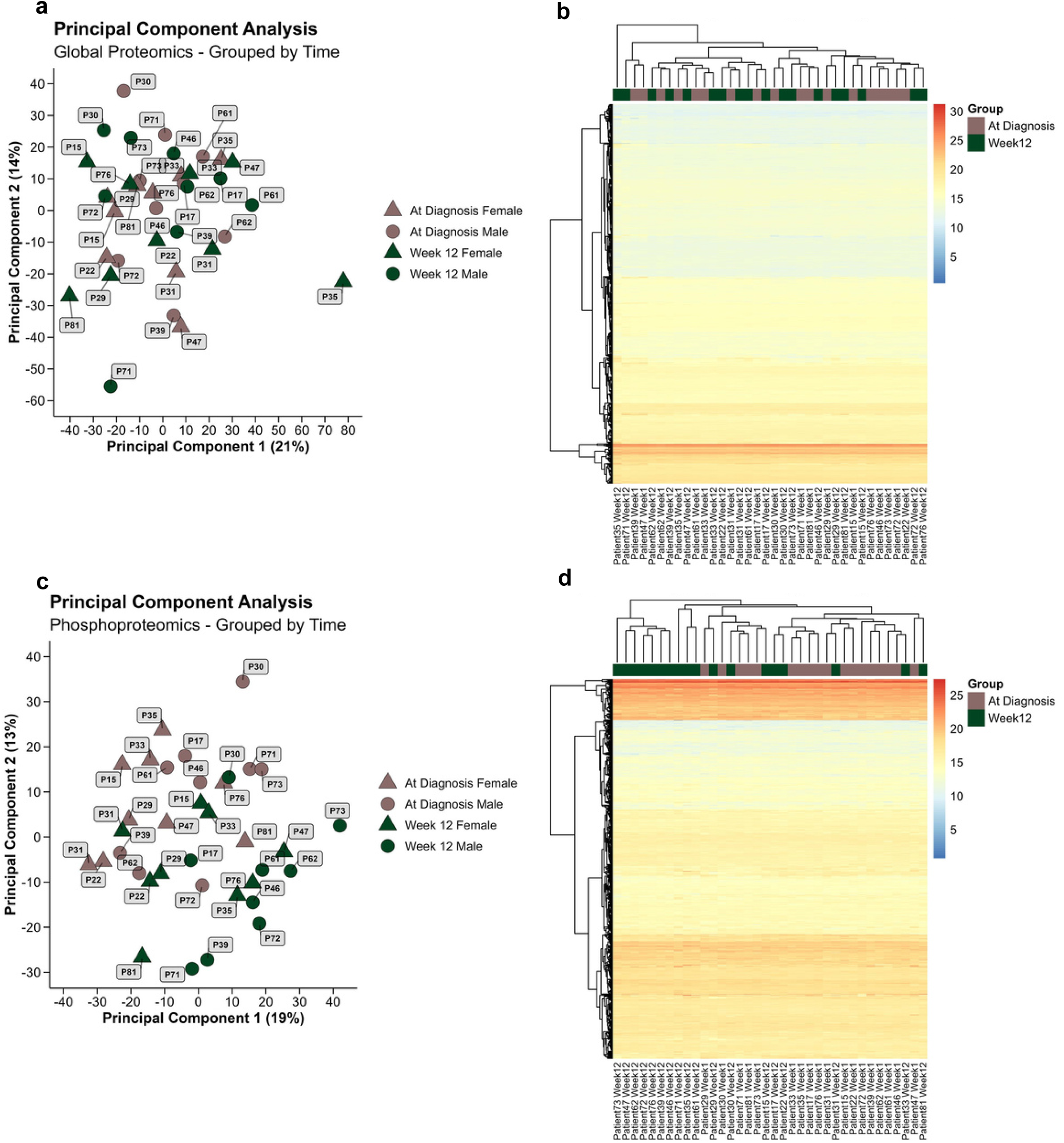
Global proteomics; paired at week 1 of diagnosis vs. week 12 **A:** Principal component analysis of protein quantified in the global proteomics analysis showing sample clustering across all patients at week 1 of diagnosis and week 12. **B:** Hierarchical clustering of protein quantified in the global proteomics analysis based on log2-transformed intensities in all patients at diagnosis and week 12. **C:** Principal component analysis of protein quantified in the phosphoproteomics analysis showing sample clustering across all patients at diagnosis and week 12. **D:** Hierarchical clustering of protein quantified in the phosphoproteomics analysis based on log2-transformed intensities in all patients at diagnosis and week 12.

**Figure S4.**
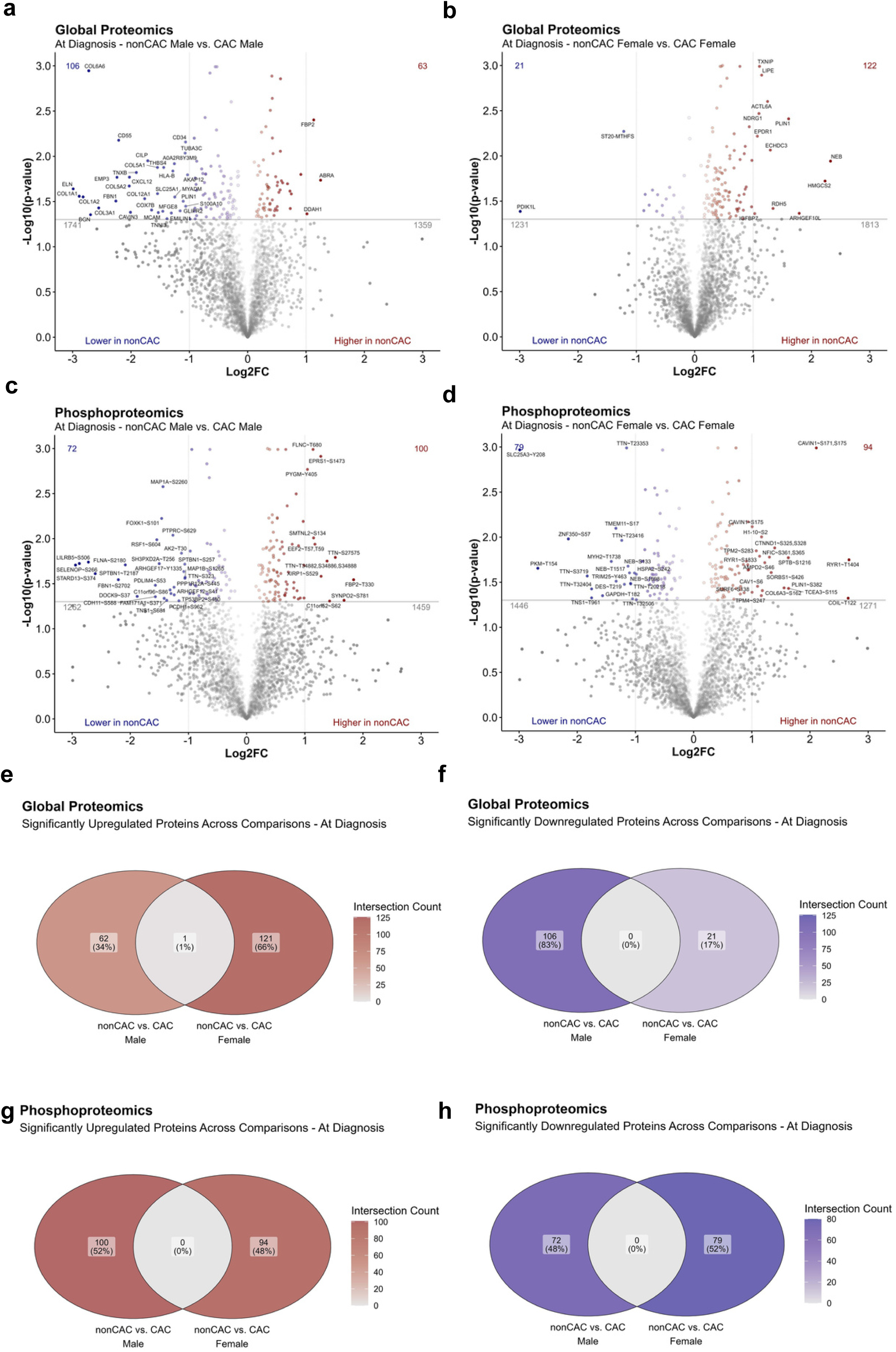
Sex-specific global and phosphoproteomic divergence associated with cachexia at diagnosis in patients with NSCLC. **A-B:** Volcano plot displaying the log2 fold change (Log2FC) and -log10 p-values of proteins quantified by global proteomics and phosphoproteomics at diagnosis between nonCAC males vs. CAC males. Significantly regulated proteins with Log2FC > 1 or < −1 are highlighted. Statistical comparisons were performed using Welch’s t-test, while adjustment for multiple-comparison was not applied. **C-D:** Volcano plot displaying the log2 fold change (Log2FC) and -log10 p-values of proteins quantified by global proteomics and phosphoproteomics at diagnosis between nonCAC females vs. CAC females. Significantly regulated proteins with Log2FC > 1 or < −1 are highlighted. Statistical comparisons were performed using Welch’s t-test, while adjustment for multiple-comparison was not applied. **E-F**: Venn diagram showing significantly overlapping up- and downregulated proteins quantified by global proteomics at diagnosis in nonCAC males vs. CAC males and nonCAC females vs. CAC females. **G-H**: Venn diagram showing significantly overlapping up- and downregulated phospho-sites quantified by phosphoproteomics at diagnosis in nonCAC males vs. CAC males and nonCAC females vs. CAC females.

**Table S1.**
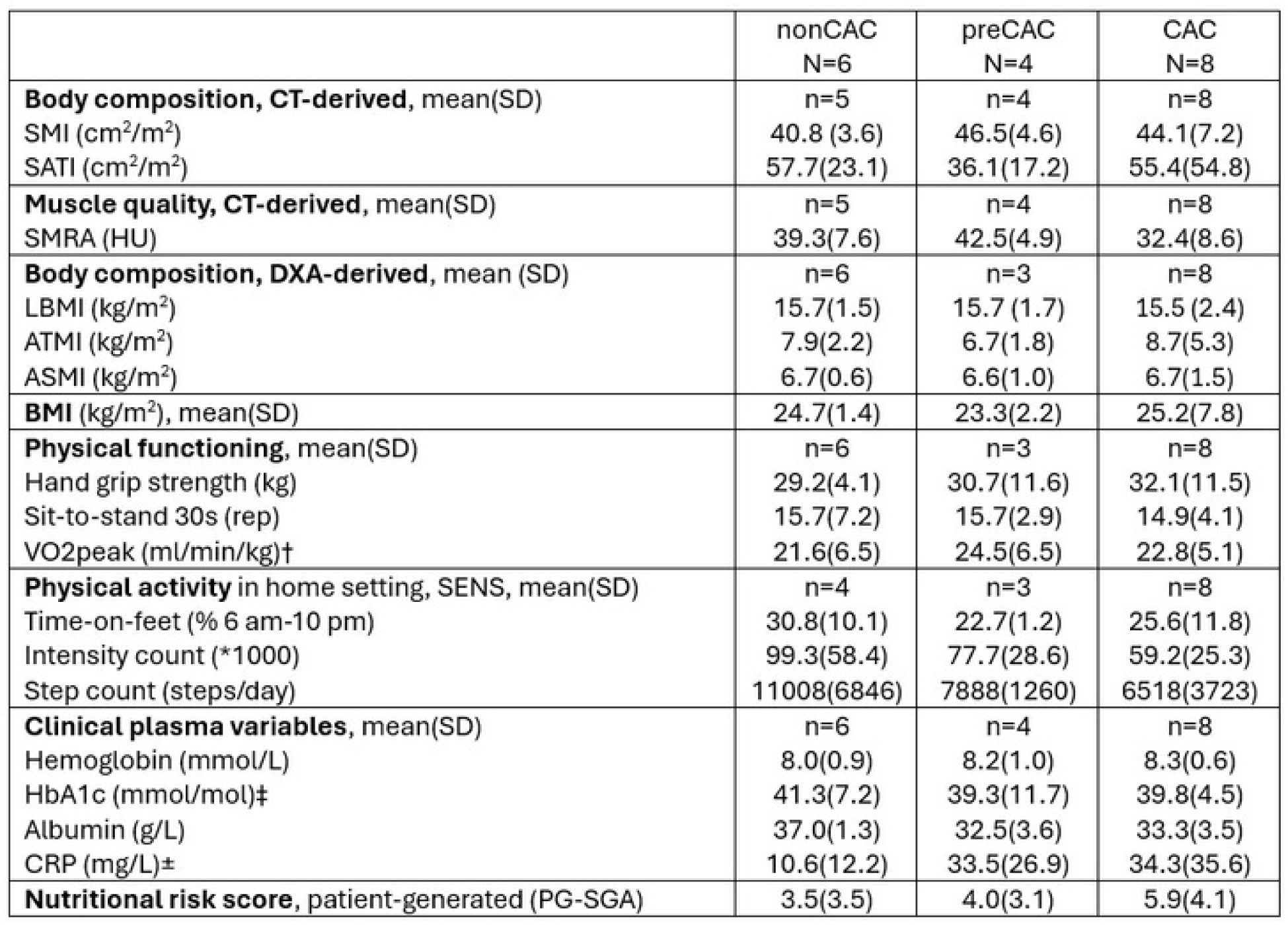
The 18 clinical variables in patients with nonCAC, preCAC and CAC at diagnosis. nonCAC=non-cachectic. preCAC=pre-cachectic. CAC=cachectic. CT=computed tomography. SD=standard deviation. cm=centimetre. m=metre. SMI=skeletal muscle index. SATI=subcutaneous adipose tissue index. SMRA=skeletal muscle radio attenuation. HU=hounsfield units. DXA=dual energy x-ray absorptiometry. LBMI=lean body mass index. ATMI=adipose tissue mass index. ASMI=appendicular skeletal muscle index. BMI=body mass index. CRP=c-reactive protein. PG-SGA=patient-generated subjective global assessment. N=numbers of patients in each group. n=numbers of patients with data for each variable. For measures not completed by all participants, sample sizes are indicated as follows: † nonCAC n=5, preCAC n=2, CAC n=6; ‡ nonCAC n=4, preCAC n=4, CAC n=5; ± nonCAC n=6, preCAC n=4, CAC n=6. One-way ANOVA comparing the three groups revealed no significant differences.

|  | nonCAC<br>N=6 | preCAC<br>N=4 | CAC<br>N=8 |
| --- | --- | --- | --- |
| **Body composition, CT-derived, mean(SD)** | n=5 | n=4 | n=8 |
| SMI (cm<sup>2</sup>/m<sup>2</sup>) | 40.8 (3.6) | 46.5(4.6) | 44.1(7.2) |
| SATI (cm<sup>2</sup>/m<sup>2</sup>) | 57.7(23.1) | 36.1(17.2) | 55.4(54.8) |
| **Muscle quality, CT-derived, mean(SD)** | n=5 | n=4 | n=8 |
| SMRA (HU) | 39.3(7.6) | 42.5(4.9) | 32.4(8.6) |
| **Body composition, DXA-derived, mean (SD)** | n=6 | n=3 | n=8 |
| LBMI (kg/m<sup>2</sup>) | 15.7(1.5) | 15.7 (1.7) | 15.5 (2.4) |
| ATMI (kg/m<sup>2</sup>) | 7.9(2.2) | 6.7(1.8) | 8.7(5.3) |
| ASMI (kg/m<sup>2</sup>) | 6.7(0.6) | 6.6(1.0) | 6.7(1.5) |
| **BMI (kg/m<sup>2</sup>), mean(SD)** | 24.7(1.4) | 23.3(2.2) | 25.2(7.8) |
| **Physical functioning, mean(SD)** | n=6 | n=3 | n=8 |
| Hand grip strength (kg) | 29.2(4.1) | 30.7(11.6) | 32.1(11.5) |
| Sit-to-stand 30s (rep) | 15.7(7.2) | 15.7(2.9) | 14.9(4.1) |
| VO2peak (ml/min/kg)† | 21.6(6.5) | 24.5(6.5) | 22.8(5.1) |
| **Physical activity in home setting, SENS, mean(SD)** | n=4 | n=3 | n=8 |
| Time-on-feet (% 6 am-10 pm) | 30.8(10.1) | 22.7(1.2) | 25.6(11.8) |
| Intensity count (*1000) | 99.3(58.4) | 77.7(28.6) | 59.2(25.3) |
| Step count (steps/day) | 11008(6846) | 7888(1260) | 6518(3723) |
| **Clinical plasma variables, mean(SD)** | n=6 | n=4 | n=8 |
| Hemoglobin (mmol/L) | 8.0(0.9) | 8.2(1.0) | 8.3(0.6) |
| HbA1c (mmol/mol)‡ | 41.3(7.2) | 39.3(11.7) | 39.8(4.5) |
| Albumin (g/L) | 37.0(1.3) | 32.5(3.6) | 33.3(3.5) |
| CRP (mg/L)± | 10.6(12.2) | 33.5(26.9) | 34.3(35.6) |
| **Nutritional risk score, patient-generated (PG-SGA)** | 3.5(3.5) | 4.0(3.1) | 5.9(4.1) |

